# Chromosomal instability by mutations in a novel specificity factor of the minor spliceosome

**DOI:** 10.1101/2020.08.06.239418

**Authors:** Bas de Wolf, Ali Oghabian, Maureen V. Akinyi, Sandra Hanks, Eelco C. Tromer, Jolien van Hooff, Lisa van Voorthuijsen, Laura E. van Rooijen, Jens Verbeeren, Esther C.H. Uijttewaal, Marijke P.A. Baltissen, Shawn Yost, Philippe Piloquet, Michiel Vermeulen, Berend Snel, Bertrand Isidor, Nazneen Rahman, Mikko J. Frilander, Geert J.P.L. Kops

## Abstract

Aneuploidy is the leading cause of miscarriage and congenital birth defects, and a hallmark of cancer. Despite this strong association with human disease, the genetic causes of aneuploidy remain largely unknown. Through exome sequencing of patients with constitutional mosaic aneuploidy, we identified biallelic truncating mutations in *CENATAC* (*CCDC84*). We show that CENATAC is a novel component of the minor (U12-dependent) spliceosome that promotes splicing of a specific, rare minor intron subtype. This subtype is characterized by AT-AN splice sites and relatively high basal levels of intron retention. CENATAC depletion or expression of disease mutants resulted in excessive retention of AT-AN minor introns in ~100 genes enriched for nucleocytoplasmic transport and cell cycle regulators, and caused chromosome segregation errors. Our findings reveal selectivity in minor intron splicing with a specific impact on the chromosome segregation process, and show how defects herein can cause constitutional aneuploidy.

## Introduction

Chromosome segregation errors in mitosis or meiosis lead to aneuploidy, a karyotype that deviates from an exact multiple of the haploid set of chromosomes. Aneuploidy is the leading cause of congenital birth defects and associated with approximately 35% of all spontaneous human abortions^1^. Furthermore, roughly 70% of human tumors are aneuploid, making it one of the most common genomic alterations in cancer^2,3^. Despite this common association of aneuploidy with human disease, little is known about its genetic causes. The study of aneuploidy-associated hereditary disorders can be instrumental in uncovering these causes.

Mosaic variegated aneuploidy (MVA; OMIM: 257300) is a rare autosomal recessive disorder characterized by mosaic aneuploidies in multiple tissues. Patients often present with microcephaly, developmental delay, various congenital abnormalities, and childhood cancers^4^. Pathogenic mutations in *BUB1B*, *CEP57*, or *TRIP13* have been identified in roughly half of all MVA patients^5–8^. These genes have well-documented roles in chromosome segregation^9–12^. All three gene products (BUBR1, CEP57 and TRIP13) promote spindle assembly checkpoint (SAC) function^10,13–16^, and BUBR1 and CEP57 additionally ensure correct kinetochore-microtubule attachment^12,17^. As predicted, such mitotic processes are defective in cells from MVA patients carrying biallelic mutations in these genes, explaining the chromosomal instability (CIN) phenotype and resulting aneuploid karyotypes. CIN can also result from mutations in regulators of expression of mitotic genes. For example, mutations in the retinoblastoma gene (*RB1*) causes CIN by over expression of the SAC protein MAD2^18–20^. In this work, we show that chromosome segregation errors can be caused by a specific defect in minor intron splicing, another process governing correct gene expression.

While the conventional, major spliceosome targets most (>99.5%) human introns, the minor spliceosome recognizes and excises only a small subset (~700 introns)^21,22^. These minor introns (also called U12-type introns) have highly conserved 5’ splice site (5’ss) and branch point (BPS) sequences that are longer and differ at the sequence level from the respective sequences in major (U2-type) introns. Most minor introns have AT-AC or GT-AG terminal dinucleotides (24% and 69%, respectively)^21,23^. In addition, the 3’ terminal nucleotide can vary, thus giving rise to AT-AN and GT-AN classes of minor introns^24,25^. For simplicity, we refer these as A- and G-type introns, respectively. Thus far there has been no indication of mechanistic or functional differences between the minor intron subtypes.

Minor intron “host” genes, the position of the minor intron within the gene, and intron subtypes are all evolutionarily conserved^21,23,26–28^. Despite this high conservation, the functional significance of minor introns has remained elusive. Elevated levels of unspliced minor introns in various cell types have been reported, giving rise to the hypothesis that these are rate-limiting controls for the expression of their host genes^29–32^. Neverteless, the overall significance of the elevated intron retention levels has been questioned particularly at individual gene level^33^.

The overall architecture of the minor and major spliceosomes is highly similar. Both are composed of five small ribonucleoprotein (snRNP) complexes containing small nuclear RNA (snRNA) molecules and a large number of proteins components. One of the snRNAs (U5) is shared between the spliceosome, while U1, U2, U4 and U6 snRNAs are specific to major spliceosome and U11, U12, U4atac, and U6atac snRNAs to the minor spliceosome. Introns are initially recognized by the U1 and U2 snRNPs (major spliceosome) or by the U11/U12 di-snRNP (minor spliceosome), followed by the entry of the U4/U6.U5 snRNP or U4atac/U6atac.U5 tri-snRNP and subsequent architectural changes leading to catalytic activation of the spliceosome^22^. At the protein level, the main difference between the spliceosomes is in the composition of U11/U12 di-snRNP that contains seven unique protein components that are needed for recognition of the unique minor intron splice sequences^34^. In contrast, the protein composition between the minor and major tri-snRNPs appears similar, but rigorous comparative analyses have been difficult due to ~100-fold lower cellular abundance of the minor tri-snRNP^35^.

Here we report that germline mutations in a novel component of the minor spliceosome (CENATAC/CCDC84) cause chromosomal instability in MVA patients. We identify CENATAC as a minor spliceosome-specific tri-snRNP subunit that selectively promotes splicing of the A-type minor introns. CENATAC mutant patient cells show high levels of A-type intron retention in only a small set of genes including ones involved in chromosome segregation, suggesting a molecular explanation for the MVA phenotype in these patients.

## Results

### Biallelic truncating mutations in *CENATAC* (*CCDC84*) cause MVA

To search for additional causes of MVA, we performed exome sequencing and variant analyses on MVA patients and family members, as previously described^8^. We identified biallelic truncating mutations in coiled-coil domain-containing 84 (*CCDC84,* hereafter named *CENATAC*, for centrosomal AT-AC splicing factor, see below) in two affected siblings with 7.3% and 8.5% aneuploid blood cells, respectively (Figs. 1A and S1). Both siblings were alive at 47 and 33 years of age and had microcephaly, mild developmental delay, and mild maculopathy. Neither individual had short stature, dysmorphism or cancer. Each parent was heterozygous for one of the mutations and the unaffected sibling had neither mutation. Moreover, the mutations were absent from the ExAC and ICR1000 series and we estimated the chance of an individual having two truncating *CENATAC* mutations to be 4.8×10-10^36^. We therefore consider it very likely that the *CENATAC* mutations are the cause of the siblings’ phenotype. The paternal and maternal mutations (mutation 1 and mutation 2, respectively) both result in the creation of novel splice sites that lead to a frameshift and the loss of the C-terminal 64 amino acids of CENATAC (Figs. 1B and S2). Although expression of the mutant alleles was very low in the parental cells, expression of the maternal allele was elevated in the cells of patient 1 (hereafter called patient), and was responsible for the low expression of wild-type protein in these cells due to infrequent recognition of the original splice site (Figs. 1C and S2C).

**Fig. 1.**
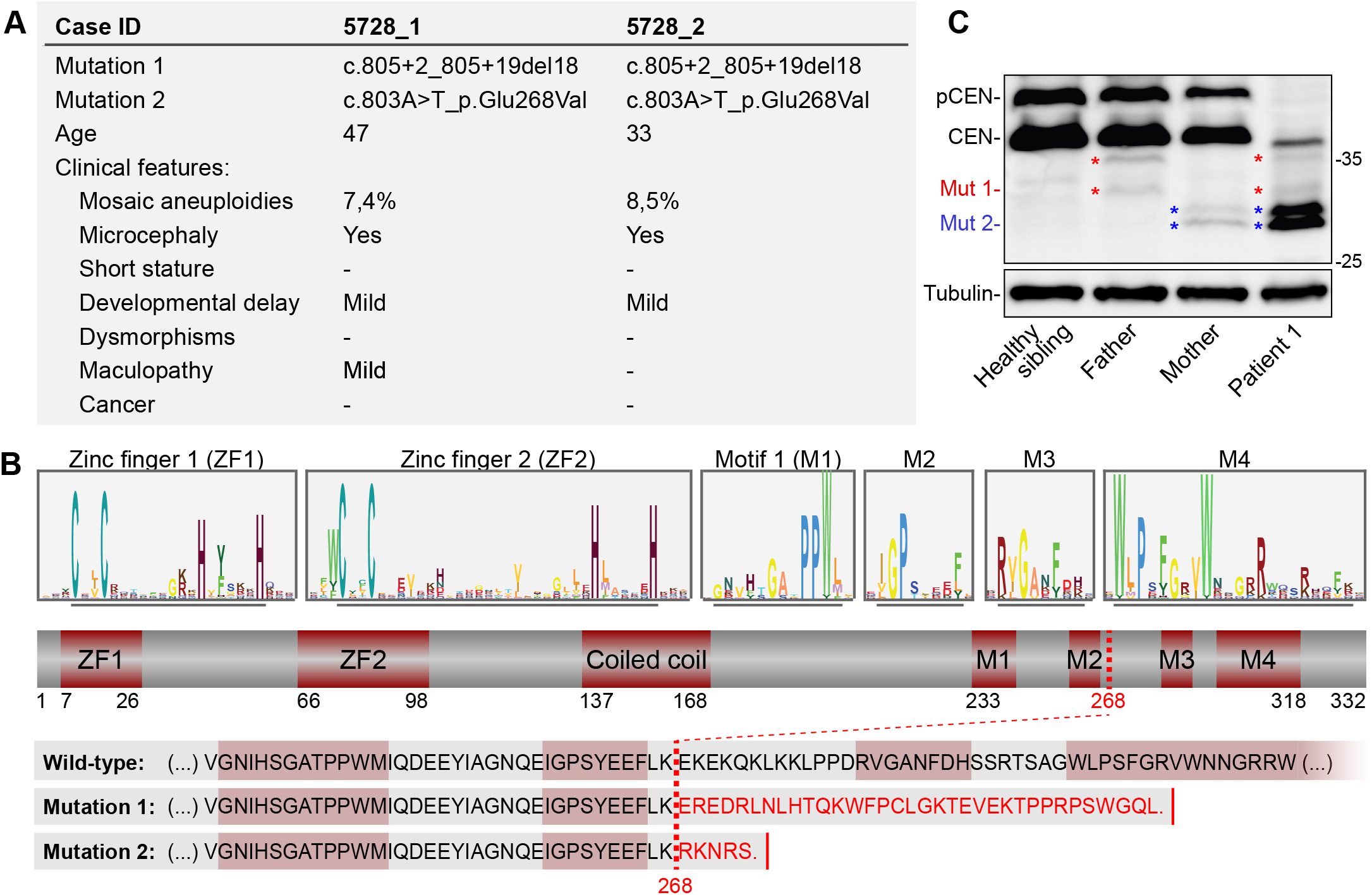
Biallelic truncating mutations in *CENATAC* (*CCDC84*) cause MVA. **A)** Clinical phenotypes of *CENATAC (CCDC84)* mutant patients. See also Fig. S1. **B)** Schematic representation of CENATAC annotated with zinc fingers (ZF1 and ZF2), predicted coiled coil, and conserved motifs 1-4 (M1-M4). Upper: sequence logos of both zinc fingers and the conserved residues defining motifs 1-4 (underlined). See Fig. S3 for the full-length logo. Lower: C-terminal protein sequences of wild-type and MVA mutant CENATAC. The MVA truncation site is indicated by the red dotted line; the four conserved motifs are outlined in red. **C)** CENATAC and tubulin immunoblots of lysates from lymphoblasts of patient 1 and relatives. Wild-type and truncated, mutant proteins are indicated. Wild-type CENATAC (CEN): 38 kDa, Mut1: 34.5 kDa (red asterisks), Mut2: 31.1 kDa (blue asterisks). pCEN indicates phosphorylated CENATAC^38^.

*CENATAC* is an essential gene whose product has previously been reported to interact with pre-mRNA splicing factors and to localize to centrosomes where it suppresses centriole over-duplication and spindle multipolarity^37,38^. Analysis of *CENATAC* sequence conservation in metazoan species revealed the presence of two N-terminal C2H2 zinc-fingers and four well-conserved C-terminal sequence motifs, of which the two most C-terminal are lost as a result of the patient mutations (Figs. 1B and S3).

### CENATAC promotes error-free chromosome segregation

Live imaging of chromosome segregation in *CENATAC*-mutant patient lymphoblasts stably expressing H2B-mNeon^8^ revealed a mild chromosomal instability phenotype, consistent with the modest levels of aneuploidy in blood cells of these patients (Figs. 1A and 2A). To examine if *CENATAC* patient mutations cause chromosomal instability, we expressed mutant *CENATAC* alleles in HeLa cells in which the endogenous loci were modified to express AID-degron-tagged CENATAC (EGFP-AID-CENATAC, Fig. S4)^39^. Efficient depletion of CENATAC through a combination of siRNA treatment and auxin addition caused chromosome congression defects and a subsequent mitotic arrest (Figs. 2B-C and S4). This phenotype was fully rescued upon re-expression of wild-type but not MVA-mutant CENATAC (Figs. 2C, S5 and S6). Similarly, CENATAC alleles missing either of the two most C-terminal conserved motifs (Fig. 1B, motifs 3 and 4) that are absent from MVA-mutant CENATAC did not rescue the mitotic defects. Instead, expression of the MVA or motif 3/4 mutants exacerbated the phenotype, suggesting that these proteins dominantly repressed the function of any residual wild-type protein (Fig. 2C). Mutations in the zinc fingers or deletion of motifs 1 or 2 only partly compromised CENATAC function (Fig. 1B and 2C).

**Fig. 2.**
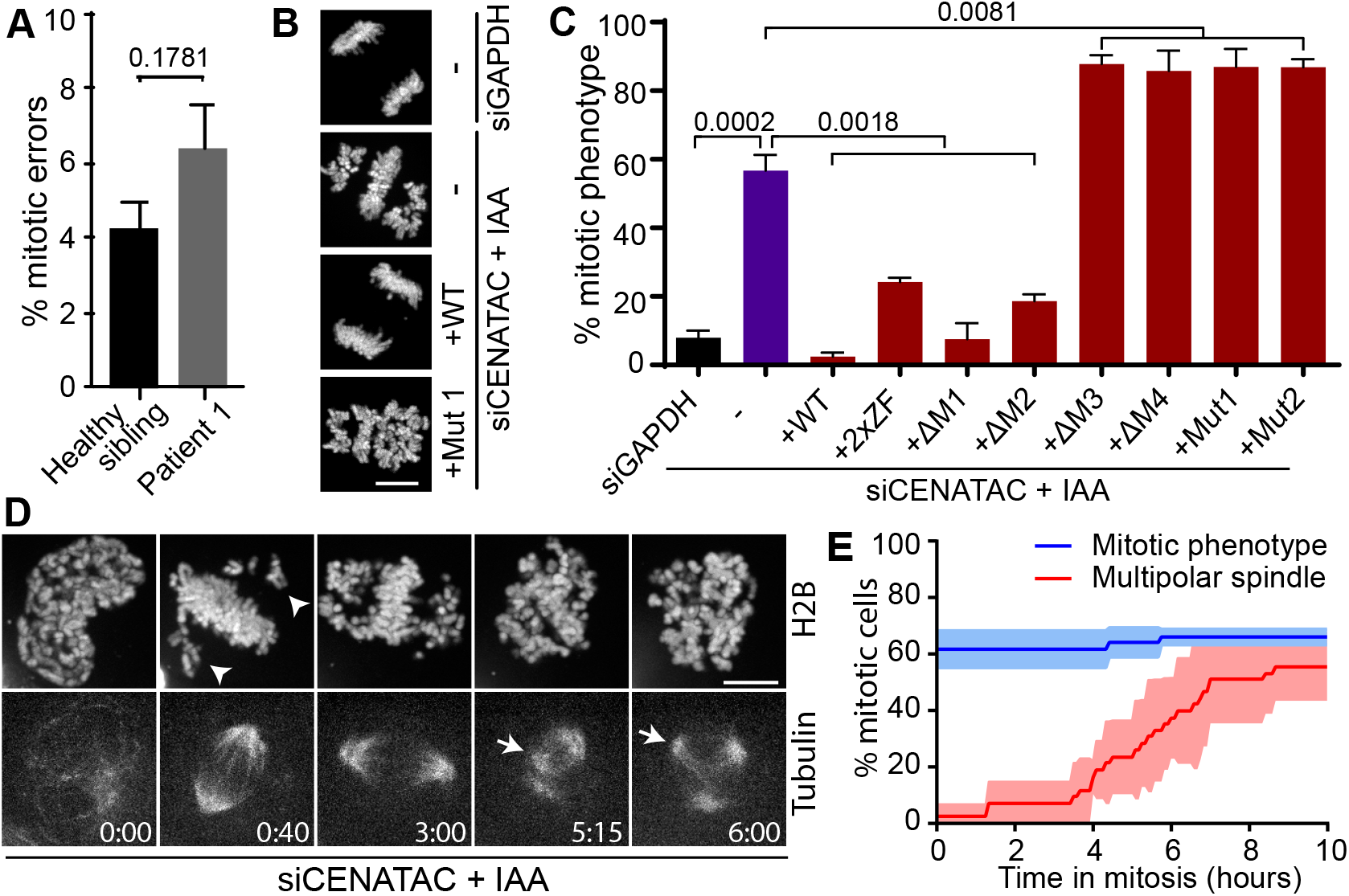
CENATAC promotes error-free chromosome segregation. **A)** Quantification of chromosome segregation errors of patient and control lymphoblasts expressing H2B-mNeon. Each bar depicts the mean of four independent experiments ± s.e.m., with >200 cells in total per condition. The P-value was calculated with a two-sided unpaired Student’s t test. **B)** Representative images of H2B-mNeon-expressing EGFP-AID-CENATAC HeLa cells depleted of GAPDH (upper) or CENATAC (middle and lower) with or without re-expression of CENATAC variants as indicated. Scale bar, 10 μm. **C)** Quantification of mitotic defects as in B) of cells treated as indicated. For 2xZF the four zinc-finger cysteines were mutated to alanines; for Δ1-4 the corresponding motif was removed. Each bar depicts the mean of three or five independent experiments ± s.e.m., with >85 cells in total per condition. P values were calculated with two-sided unpaired Student’s t tests. **D)** Representative stills of EGFP-AID-CENATAC HeLa cells expressing H2B-mNeon and depleted of CENATAC. Microtubules were visualized with SiR-Tubulin. Arrowheads and arrows indicate non-congressed chromosomes and supernumerary spindle poles, respectively. Scale bar, 5 μm. Time in hours. See Fig. S7 for the control condition. See also Movies S1 and S2. **E)** Quantification of the mitotic phenotype and multipolar spindle formation in time in cells treated as in E). Each line depicts the mean of three experiments ± s.e.m., with >44 cells in total per condition. See also Fig. S7.

Live imaging of EGFP-AID-CENATAC cells with fluorescently labelled chromatin and microtubules revealed that the chromosome congression defect upon CENATAC depletion preceded the previously described loss of spindle bipolarity (Figs. 2D-E and S7, Movies S1 and S2)^38^. In addition, we did not observe centriole over-duplication in CENATAC-depleted cells (Figs. S7C-D). This is in contrast to what was recently reported for *CENATAC* knockout cells^38^, raising the possibility that centriole over-duplication is a cumulative effect of prolonged CENATAC loss. Our attempts to examine this failed, as we were unable to create *CENATAC* knockout cells, consistent with it being an essential human gene^37,40,41^. Taken together, these data show that CENATAC has a function in promoting chromosome congression in mitosis, which is likely unrelated to its role in maintaining spindle bipolarity, and that MVA mutant CENATAC is defective for this function.

### CENATAC is a novel component of the minor spliceosome

To investigate in which processes CENATAC plays a role, we performed a genome-wide, evolutionary co-occurrence analysis. Genes that function in the same biochemical process experience similar evolutionary pressures and therefore tend to co-evolve, i.e. they are lost or retained in a coherent fashion^42^. Genomes from a set of 90 informative eukaryotic species (Table S1) were mined for the presence or absence of *CENATAC* orthologs^43^. This provided a phylogenetic absence/presence profile that was used in an unbiased genome-wide query for genes with similar phylogenetic profiles (Fig. 3A). The resulting list of genes most strongly co-occurring with *CENATAC* was significantly enriched for components of the minor (U12-dependent) spliceosome complex (Fig. 3B, Table S2). We thus reasoned that CENATAC may play a role in splicing by the minor spliceosome.

**Fig. 3.**
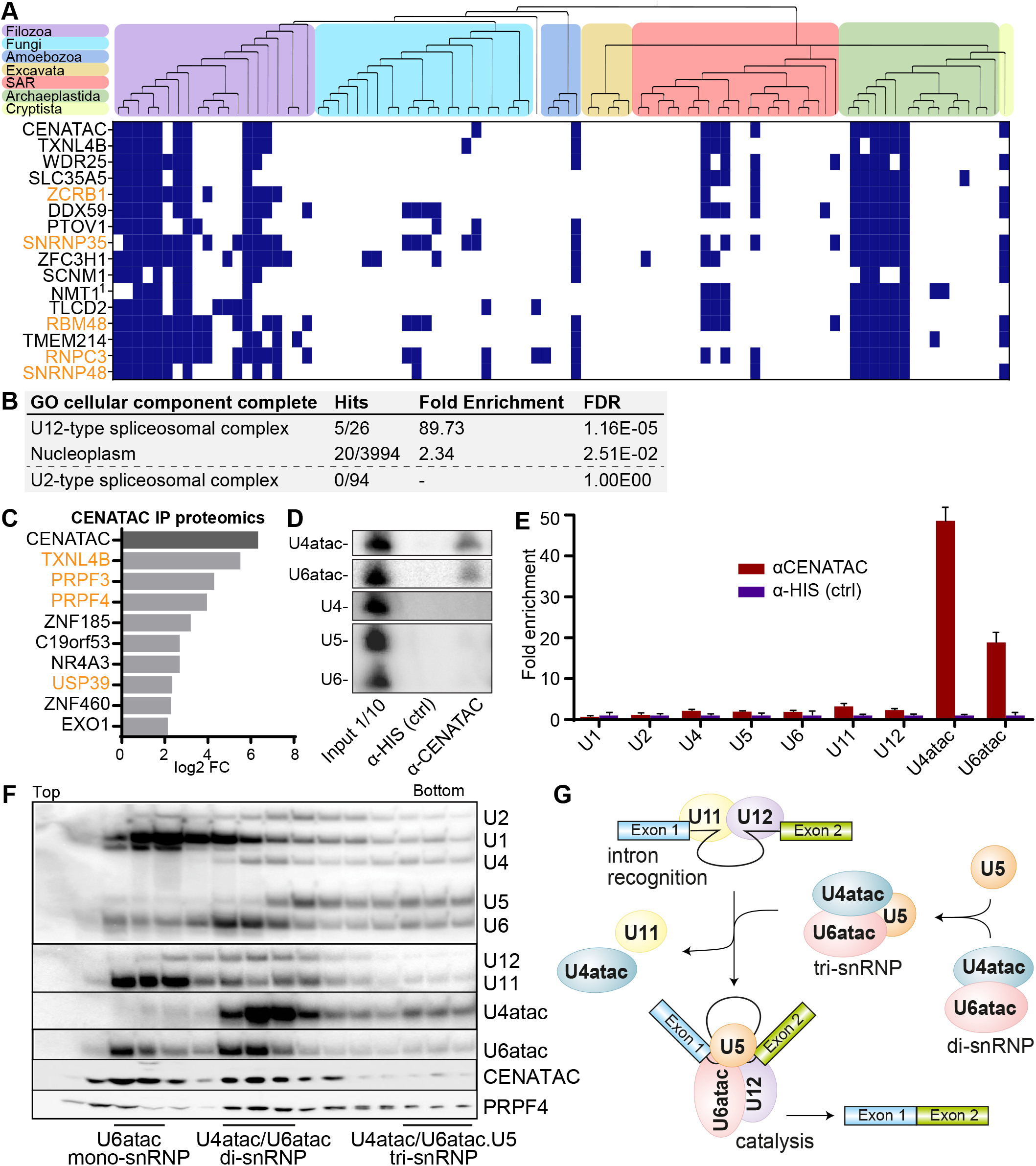
CENATAC is a novel component of the minor spliceosome. **A)** Phylogenetic profiles (presences (blue) and absences (white)) of the top 15 genes co-occurring with *CENATAC* in 90 eukaryotic species. Top: phylogenetic tree of the eukaryotic species (see Table S1) with colored areas for the eukaryotic supergroups. ^1^ for *NMT1* no human ortholog was found and instead the *Arabidopsis thaliana* ortholog is depicted. Genes associated with the minor (U12-dependent) spliceosome are depicted in orange. See also Table S2. **B)** GO term analysis of the genes co-occurring with *CENATAC* as in A) with a correlation score of >0.5. **C)** Graph of fold-changes of proteins enriched (P-value < 0.05) in proteomics analysis of CENATAC vs. control co-immunoprecipitations. Each bar depicts the fold change of the mean log 2 transformed LFQ intensity of three independent experiments. Splicing factors are depicted in orange. See also Fig. S8. **D-E)** Examples **(D)** and quantification **(E)** of Northern blot analyses of minor (U6atac, U4atac, U11, and U12) and major (U2, U1, U4, U5, and U6) spliceosome snRNAs. Each bar depicts the mean of 3 experiments normalized to the control ± std. dev. See also Fig. S9. **F)** Glycerol gradient analysis of HeLa nuclear extracts. snRNAs were detected by Northern blot analysis, proteins (CENATAC and PRPF4) by western blot. Locations of the U6atac mono-snRNP, U4atac/U6atac di-snRNP and U4atac/U6atac.U5 tri-snRNP are indicated. **G)** Schematics showing key assembly stages in minor intron splicing and minor tri-snRNP assembly: intron recognition (A-complex) and the catalytic spliceosome (C-complex). For simplicity, several stages of spliceosome assembly are omitted, such as the pre-B-complex, which consists of the the intron recognition complex together with the tri-snRNP before architectural changes lead to the exclusion of U11 to give rise to the B complex, after which subsequent architectural changes lead to the exclusion of U4atac to give rise to the B^ACT^ complex, which is a precursor stage for the catalytically active C complex depicted in this figure^48^.

As predicted by our co-evolution analysis and in agreement with a previous high-throughput screen^37,44–46^, mass spectrometry analysis of proteins co-purifying with CENATAC identified several known spliceosome components that are shared by both the major and minor spliceosomes (Figs. 3C and S8). Notably, the strongest CENATAC interactor (TXNL4B) was also the gene that showed the most significant co-occurrence with CENATAC in eukaryotic species (Fig. 3A). To determine whether CENATAC preferentially associates with major or minor spliceosome components, we analyzed CENATAC co-immunoprecipitations by Northern blot analysis. This revealed a significant enrichment for the minor spliceosome-specific U4atac and U6atac snRNAs (Figs. 3D-E and S9) CENATAC’s association with the minor spliceosome was further supported by glycerol gradient analyses of HeLa nuclear extract preparations, which showed co-migration of CENATAC with U6atac snRNP, U4atac/U6atac di-snRNP and U4atac/U6atac.U5 tri-snRNP complexes (Fig. 3F). Together, these data validate CENATAC as a bona fide functional component of the minor spliceosome and the first identified protein component that is specific to the U4atac/U6atac and U4atac/U6atac.U5 snRNP complexes (Fig. 3G).

The role of *CENATAC* in minor spliceosome function was further supported by the presence of evolutionarily conserved competing U2-type and U12-type 5’ splice sequences (5’ss) in animals that are predicted to generate productive and unproductive *CENATAC* mRNAs, respectively (Fig. S10). This configuration is indicative of an autoregulatory circuit that is conceptually similar to the previously reported autoregulation of the minor spliceosome proteins 48K and 65K^47,48^. In agreement with this, impaired minor spliceosome function, such as in Taybi-Linder syndrome (TALS/ MOPD1, OMIM: 210710) patients, leads to a significant increase in the use of U2-type 5’ss and upregulation of *CENATAC* mRNA levels^49^. Notably, evidence for a similar autoregulatory circuit is also present in plants (Fig. S10), where retention or splicing of a U12-type intron results in productive or unproductive *CENATAC* mRNA, respectively.

### Minor intron splicing defects in *CENATAC* mutant cells fully correlate with mitotic defects

In agreement with our finding that CENATAC is a novel minor spliceosome component, splicing of several minor (U12-type) introns was impaired upon depletion of CENATAC, whereas up- or downstream major (U2-type) introns were unaffected (Figs. 4A-B). This was true also for *CENATAC*-mutant MVA patient cells (Fig. 4B). Importantly, the splicing defect of U12-type introns in CENATAC-depleted cells was fully rescued by re-expression of wild-type but not MVA-mutant alleles (Fig. 4A). Similar to the mitotic phenotype, expression of the disease alleles and mutants lacking motifs 3 and 4 exacerbated the splicing defect, whereas mutations in the zinc fingers and removal of motifs 1 and 2 partially rescued it (Fig. 4A). Notably, the extent of the splicing defect strongly correlated with the extent of the mitotic phenotype for all mutations (Fig. 4C), indicating that it is likely that the chromosome congression phenotype is a secondary effect of impaired minor spliceosome function.

**Fig. 4.**
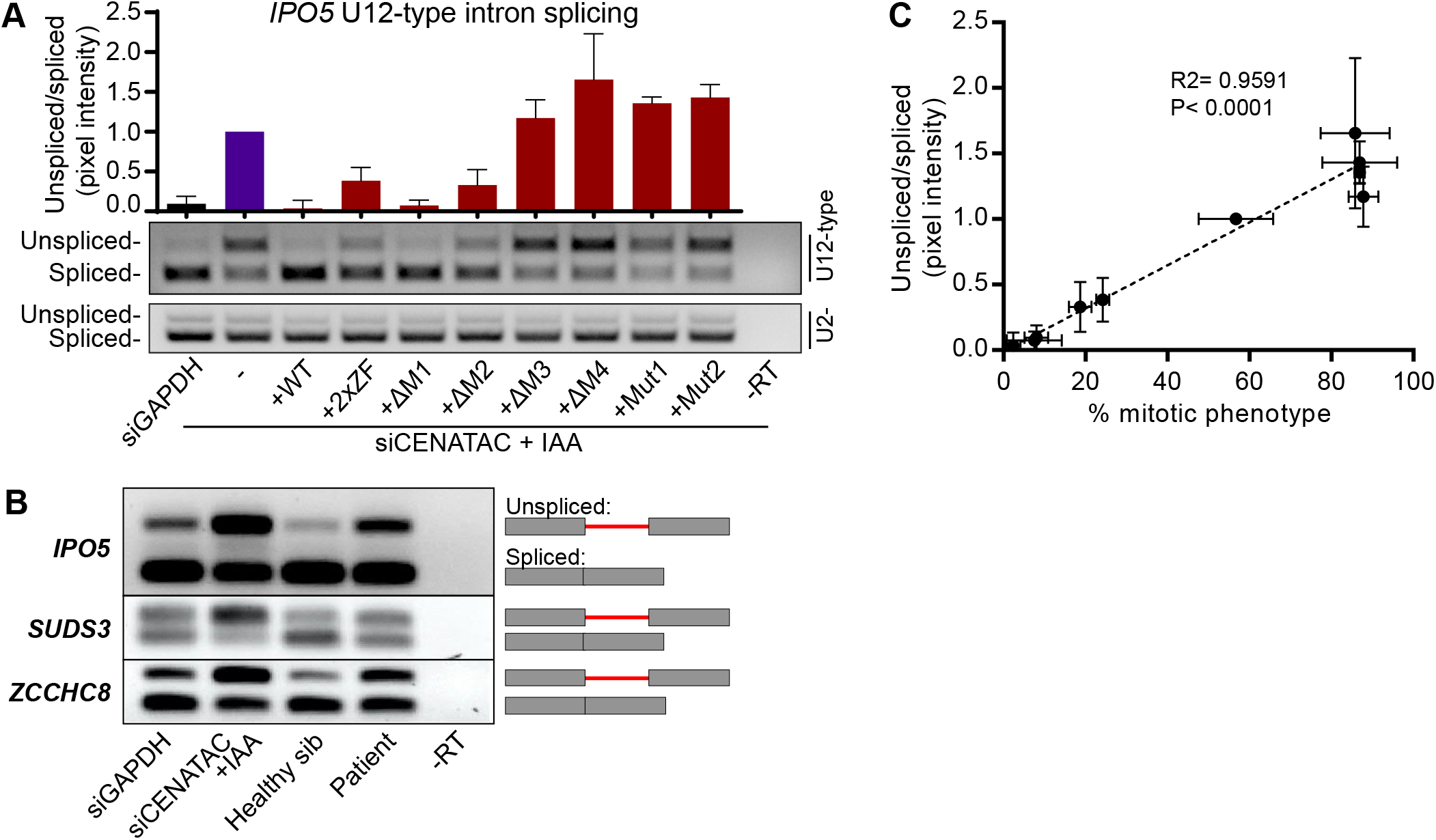
Minor intron splicing defects in *CENATAC* mutant cells fully correlate with mitotic defects. **A)** RT-PCR (middle panel) and quantification (upper panel) of *IPO5* minor intron 21 (U12-type intron) splicing on RNA extracted from cells treated as in Fig. 2C. Each bar depicts the mean of three independent experiments ± s.e.m., normalized to siCENATAC + IAA. The bottom panel shows RT-PCR analysis of IPO5 intron 19 (U2-type intron). **B)** RT-PCRs of *IPO5, SUDS3 and ZCCHC8* minor introns on RNA extracted from patient lymphoblasts or EGFP-AID-CENATAC HeLa cells depleted of GAPDH or CENATAC as indicated. **C)** For each condition in A), U12-type intron splicing (Fig. 4A) was plotted against the percentage of cells showing a mitotic phenotype (Fig. 2C). R2 and p-values are provided for the linear regression trendline (dotted line).

### CENATAC selectively promotes splicing of A-type minor introns

Our discovery of reduced minor spliceosome function in a constitutional aneuploidy syndrome raised the question of which introns and transcripts were affected by CENATAC malfunction. To investigate this, we compared the transcriptomes of CENATAC-depleted HeLa cells to those of control-depleted and parental cell lines. The resulting RNAseq dataset was analyzed for changes in intron retention (IR) using IntEREst^50^ and for alternative splicing (AS) using Whippet^51^. In agreement with our RT-PCR-based observations (Fig. 4), this analysis confirmed the significant retention of U12-but not U2-type introns after CENATAC depletion. Surprisingly, it also uncovered a remarkable enrichment for a specific subclass of U12-type introns: while only 24% of G-type introns (with GT-AG, GT-AT, GT-TG, GC-AG terminal dinucleotides) were affected by CENATAC depletion, virtually all (92%) of the A-type introns (with AT-AC^i^, AT-AA, AT-AG, or AT-AT terminal dinucleotides) showed increased retention or activation of alternative U2-type splice sites (cryptic or annotated), or both (Figs. 5A-B, S11 and Table S3). For comparison, we carried out the same analysis on a previously published dataset derived from myelodysplastic syndrome (MDS) patients carrying somatic mutations in the gene encoding for the U11/12-di-snRNP subunit *ZRSR2*^52^. This dataset showed a nearly identical response for A- and G-type introns (Fig. 5B and Table S4). Whereas depletion of CENATAC or mutations in *ZRSR2* led to an average increase of approx. 36% and 19% in retention of A-type introns, respectively (average ΔΨ_CENATAC_=~0.36 and average ΔΨ_ZRSR2_=~0.19), G-type introns were only strongly affected by *ZRSR2* mutations (average ΔΨ_CENATAC_=~0.07 and average ΔΨ_ZRSR2_=~0.19, Figs. 5C-D, and S12). Importantly, the effect on A-type introns was specific to U12-type introns as none of the 85 major AT-AC introns or related subtypes responded to CENATAC depletion (Figs. 5Ε and Table S3). The same subtype-specific effect on minor intron splicing was also observed in *CENATAC*-mutant MVA patient lymphoblasts (Figs. 5F, S13 and Table S5), in which the affected introns correlated strongly with those affected by CENATAC depletion (Figs. 5G and S13B). Notably, the strongly affected transcripts did not include any of the genes known to cause MVA but did contain various mitotic regulators (Figs. S14 and S15, Table S3). We were unable to rescue mitotic defects by restoring expression of individual candidate targets (*CEP170*, *SPC24*, *SMC3*, and *DIAPH2*, data not shown) in CENATAC-depleted cells, suggesting that correct splicing of multiple such transcripts may be required for promoting efficient chromosome congression.

**Fig. 5.**
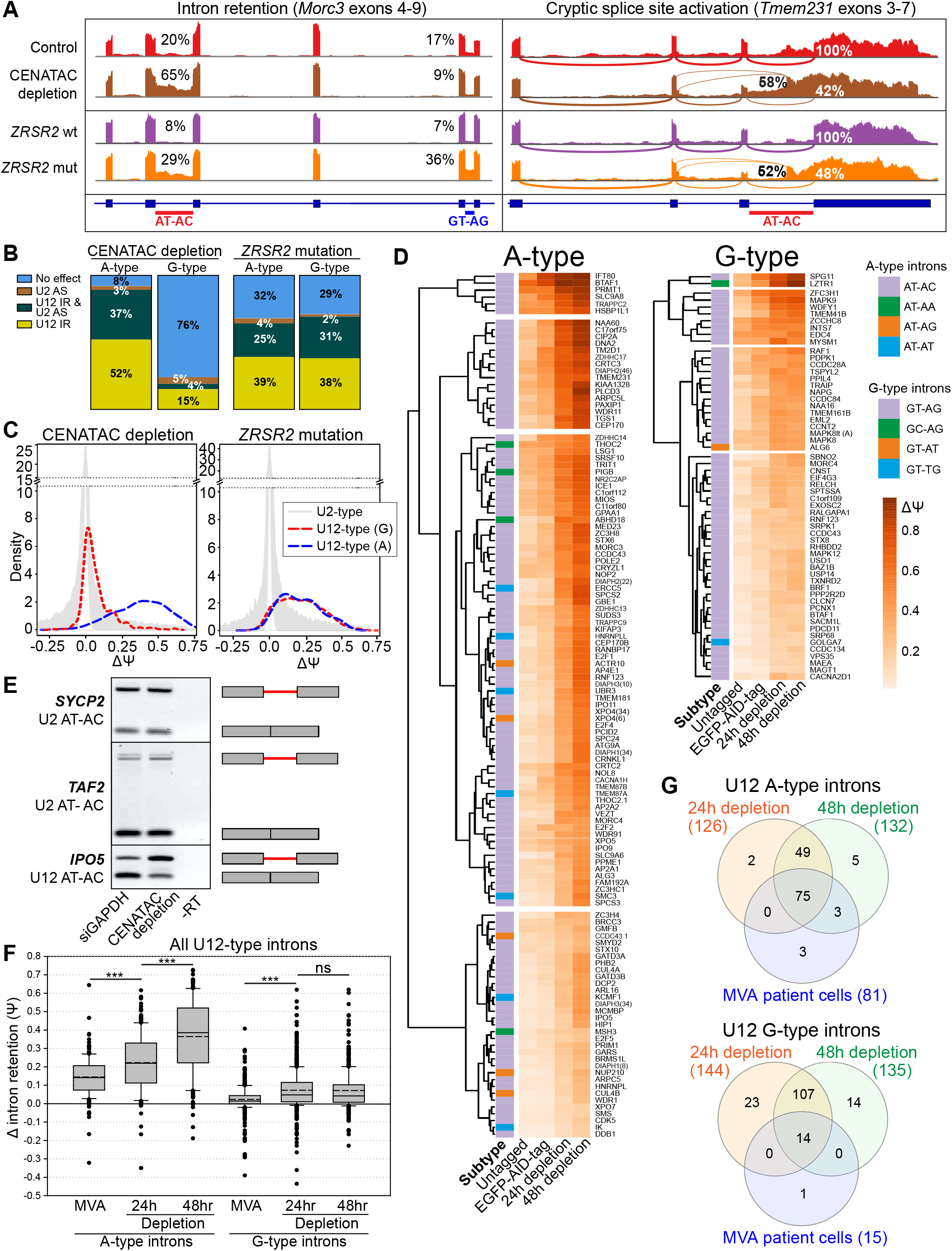
CENATAC selectively promotes splicing of A-type minor introns. **A)** Sashimi plots showing the effect of CENATAC depletion (48hr) or *ZRSR2* mutations on AT-AC and GT-AG intron retention (*Morc3*, left panel ), and AT-AC intron retention and cryptic splice site activation (*Tmem231*, right panel; percentages depict relative wild-type or cryptic splice site usage). CENATAC control represents the parental unedited HeLa cell line. **B)** Transcriptome-wide statistics of CENATAC depletion (48hrs) and *ZRSR2* mutations on G- and A-type minor intron retention (U12 IR) and cryptic U2-type splice site activation (U2 AS). Only introns showing at least 5 exon-exon junctions reads were included. For U12 IR, a statistical cutoff of Padj <0.05 was used. For U2 AS the probability cutoff of Pr > 0.9 was used. **C)** Density plots showing differences in intron retention (ΔΨ) distribution after CENATAC depletion (48 hours) or in samples with *ZRSR2* mutations. **D)** Hierarchical clustering of A- and G-type intron retention in the unedited parental cell line treated with siGADPH for 48h or in the EGFP-AID-CENATAC cell line treated with siGADPH for 48h or auxin and siCENATAC for 24 or 48 hrs. Only introns showing a Padj< 0.05 and ΔΨ>0.1 in either the 24hr or 48hr depletion sample were included in the analysis. The A-type and G-type intron terminal dinucleotide subtypes are indicated with different colors in the first column. In case the gene contained multiple introns of the same type, the intron number is indicated in parentheses. **E)** RT-PCR of U2-type AT-AC introns in *SYCP2* (intron 5) and *TAF2* (intron 1), and U12-type AT-AC intron in *IPO5* (intron 21) on RNA extracted from cells depleted of GAPDH or CENATAC. Schematic representations of unspliced/spliced PCR products are depicted on the right. **F)**Δ intron retention values for the MVA patient dataset (compared to the healthy sibling) using all (significant and not significant) U12 A-type introns (n=179 for the depletion and n=177 for the MVA patient datasets) and U12 G-type introns (n=441 for the depletion and n=446 for the MVA patient datasets). Only introns with on average at least 5 intron-mapping reads were used in the analysis. The boundaries of the boxes indicate 25th and 75th percentiles. Whiskers indicate the 90th and 10th percentiles. Median is indicated with solid line, mean with dashed line inside the box. *** - P<0.001, ns - P>0.05 (Mann-Whitney Rank Sum Test). See also Fig. S13. **G)** Venn-diagram analysis of the MVA and CENATAC depletion datasets. A- and G-type minor introns showing statistically significant intron retention in each dataset were used. See also Table S6.

### A-type minor introns are spliced less efficiently

We next wished to understand the selectivity of CENATAC-dependent splicing for A-type minor introns. The observation that also some G-type introns were affected by CENATAC depletion (Fig. 5D) argued against a direct interaction between CENATAC and intron-terminal nucleotides. Moreover, A- and G-type introns that were strongly affected by CENATAC depletion had higher basal levels of intron retention in control conditions compared to those unaffected by CENATAC depletion (Fig. 6A). This suggested that CENATAC predominantly promotes splicing of U12-type introns that are normally spliced less efficiently, and that A-type introns as a group belong to this category. To test this hypothesis we engineered the widely-used P120 minigene^53^ to contain two tandem competing A- or G-type splice sites in all possible configurations (Fig. 6B). Significantly, U12-type GT-AG splice sites were strongly preferred over AT-AC sites when in direct competition (Figs. 6B-C), and they also outcompeted the unique GC-AG splice site that was significantly affected by CENATAC depletion (Figs. 5D, S16, *LZTR1*). We thus conclude that CENATAC promotes splicing of minor introns that are recognized or spliced less efficiently, most prominently A-type minor introns.

**Fig 6.**
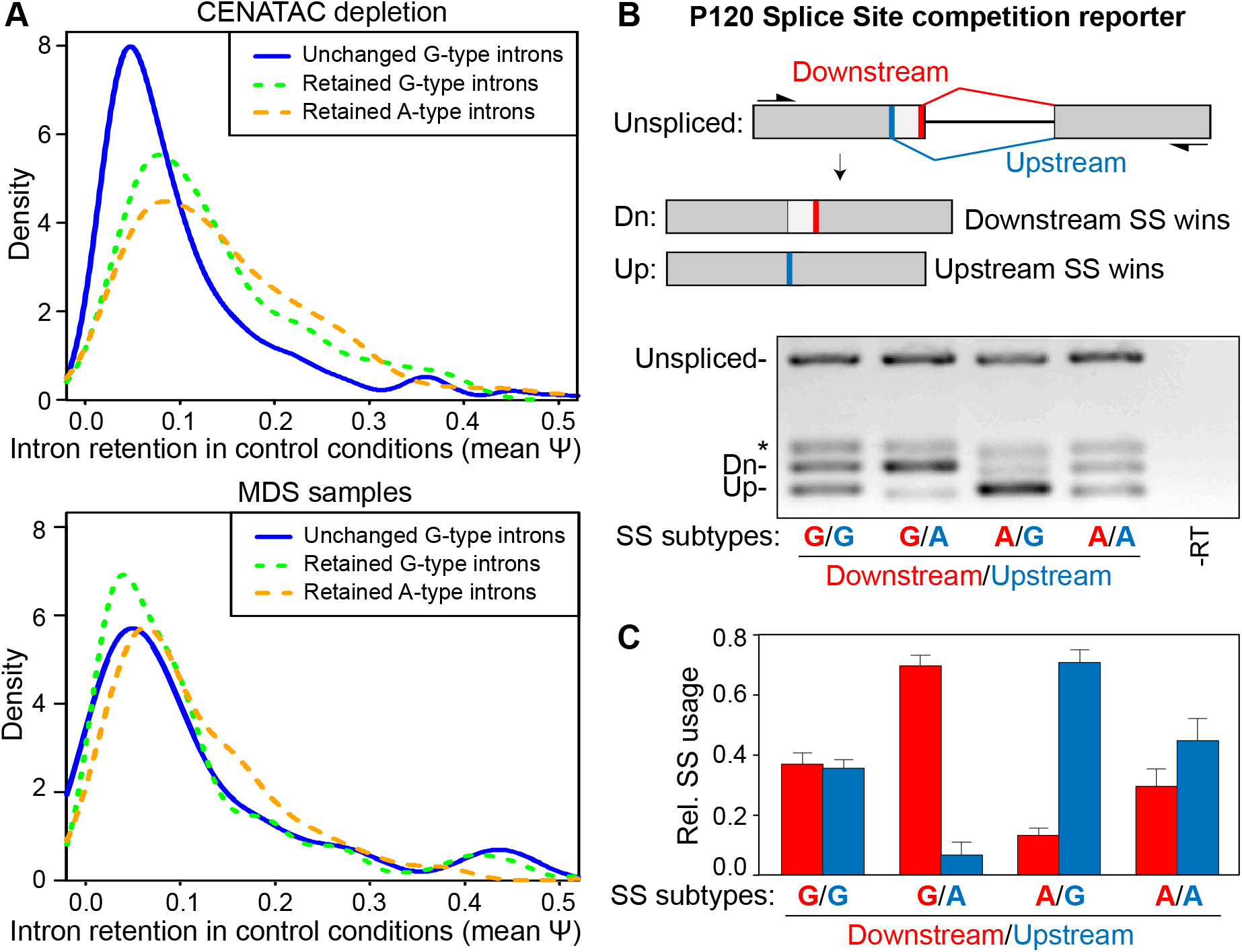
A-type minor introns are spliced less efficiently. **A)** Density plots showing intron retention (Ψ) values in the unedited parental control cell line (depleted of GAPDH) of A- and G-type minor introns that were either unchanged or retained after or CENATAC depletion (top) or *ZRSR2* mutation (bottom).The median psi-values of the retained G- and A-type introns are significantly higher (Psi=0.130 and Psi=0.115, respectively; P<0.01, Mann-Whitney Rank Sum Test) compared to the unchanged introns (Psi=0.081) in the CENATAC depletion dataset. **B)** RT-PCR P120 reporter assay^53^ to measure the relative usage of A-type (AT-AC) and G-type (GT-AG) 5’ splice sites in direct competition. Upper: schematic diagram showing the overall architecture of the reporter construct with its down- and upstream splice site (thick red and blue bars, respectively) and the products created by splicing (Dn and Up, respectively). Lower: RT-PCRs of the reporter with A- or G-type splice sites in the down- or upstream positions as indicated below the gel. SS, splice site. *, PCR product after use of a cryptic U2-type splice site (not shown in the schematic). **C)** Quantification of relative splice site usage of A- and G-type splice sites in B). Crytic splice site usage (* in B) was omitted from this plot but included in the quantification.

## Discussion

In this work, we have uncovered a novel link between the minor spliceosome and defects in chromosome segregation in human cells. Using patient exome sequencing, evolutionary co-occurrence analysis and biochemistry, we identified CENATAC as a novel protein component of the U4atac/U6atac di-snRNP and U4atac/U6atac.U5 tri-snRNP that are necessary for the formation of the catalytically active U12-dependent spliceosome. Our RNAseq analysis of both patient cells and cells depleted of CENATAC revealed widespread defects in minor intron splicing, particularly intron retention, but also cryptic splice site activation, indicating that CENATAC is important for minor spliceosome function. Unexpectedly, intron retention in CENATAC depleted cells was strongly biased for A-type minor introns, which is a subtype that is defined by AT-AN dinculeotide splice sites. This intron subtype-specific function is unique among the minor spliceosome components and correlates tightly with mitotic fidelity. Minor intron subtype missplicing is therefore likely responsible, possibly in conjunction with centriolar defects, for the inefficient chromosome congression in CENATAC-mutant cells and for the aneuploidies observed in the two MVA siblings.

The minor spliceosome was originally thought to splice only introns with AT-AC termini^53–55^. Only later it was shown that there are also U2-type AT-AC introns and that most of the minor introns in fact have GT-AG termini^56–58^ with infrequent variation in the 3’ terminal nucleotide^24,25^, thus giving rise to AT-AN and GT-AN classes of minor introns that we here referred to as A- and G-type introns. However, none of the subsequent studies, including several transcriptome-wide investigations of minor spliceosome diseases, have been able to functionally differentiate the two types of minor introns. Rather, in all known minor spliceosome diseases for which comprehensive transcriptome data is available, splicing defects are roughly uniformly distributed between the A- and G-type introns^49,52,59,60^ (see Fig. 5B). Thus, the selective A-type intron retention phenotype cannot be explained by a general loss of minor spliceosome functions but instead suggests that CENATAC has a unique, more defined function. Our observation that also a subset of G-type introns are sensitive to CENATAC depletion argues against a direct recognition of the intron 5’ adenosine by CENATAC. Instead, our competition data and the observed elevated baseline activity of the affected introns suggest that a subset of minor introns with reduced intrinsic splicing activity may be particularly dependent upon CENATAC activity. This group includes nearly all A-type minor introns. Given that A-type minor intron host genes and the locations of these introns within the host gene are evolutionarily highly conserved among metazoan species, the selectivity of CENATAC in splicing raises the possibility that minor intron subtypes are part of a conserved but unexplored regulatory mechanism for gene expression. CENATAC undergoes reversible modifications (acetylation and phosphorylation)^38^, which may provide the means to regulate its activity (also) in the minor spliceosome.

What could be the molecular function of CENATAC? Given its participation in di- and tri-snRNP complexes and sensitivity on 5’ss identity, CENATAC may function during or after the transition from initial intron recognition (A-complex) to pre-catalytic spliceosome (B-complex) (Fig. 3G) and may for instance participate in 5’ss recognition analogous to U11-48K protein in the A-complex^61^. For a deep mechanistic understanding, high-resolution structures of relevant minor spliceosome complexes are required. Nevertheless, we note that TXNL4B, the main interactor of CENATAC (Figs. 3A, 3C and S8), is a paralog of the major spliceosome protein TXNL4A/Dim1, both at sequence and structural level^62^. High-resolution structures of both yeast and human spliceosomal B-complexes have placed TXNL4A/Dim1 in close proximity of the 5’ss and suggested a role in 5’ss recognition^63,64^. Assuming that the minor spliceosome B-complex shares the molecular architecture with its major spliceosome counterpart, this would place CENATAC with TXNL4B near the 5’ss to participate in the recognition event.

Presently all mutations associated with MVA have been mapped to genes that are known regulators of chromosome segregation. In contrast, while CENATAC has been reported to regulate centriole duplication^38^, our data argue that the chromosomal instability phenotype is likely the result of a primary defect in splicing of minor introns. Our attempts to rescue the congression defect by expressing individual genes for mitotic regulator with significant A-type intron retention (*CEP170, SPC24*, *SMC3*, and *DIAPH2*) failed, suggesting that the defect may be a result of a concerted action affecting multiple proteins simultaneously. This exemplifies that chromosomal instability and the resulting aneuploid karyotypes can result from a specific defect (retention of a rare intron subtype) in a general cellular process (pre-mRNA splicing). Similarly, among the human minor spliceosomal diseases, CENATAC-linked MVA represents a unique example where the disease leads to a discrete cellular defect, in this case in chromosome segregation.

Interestingly, the clinical phenotype of CENATAC-mutant MVA resembles that of MOPD1/TALS, Roifman and Lowry-Wood syndromes, which are caused by mutations in the U4atac snRNA components of the minor spliceosome. Patients with these syndromes likewise present with microcephaly, developmental delay, and retinal abnormalities^65^. No aneuploidies were found in some studies^66,67^, but karyotype analyses were not reported for the majority of patients. It will therefore be of interest to examine if aneuploidies occur in some of these patients, and if (and to what extent) affected transcripts and the extent of the splicing defect differ between MVA and these syndromes.

## Acknowledgments

We thank the family members for their participation in this study. We thank Anna Zachariou for assistance with recruitment, Emma Ramsay for performing the exome sequencing and Elise Ruark for discussions about the analyses. We thank the Kops, Frilander, Snel and Rahman labs for discussions and comments on the manuscript. We thank Benjamin Rowland and Andrew Holland for reagents. The Kops and Vermeulen labs are part of the Oncode Institute, which is partly funded by the Dutch Cancer Society. This study was further funded by the Cancer Genomics Center (CGC.nl), the Wellcome Trust (100210/Z/12/Z) to NR, Sigrid Jusélius Foundation (MF), Jane and Aatos Erkko Foundation (MF), Academy of Finland grant 1308657 (MF), and a Postdoctoral Research Fellowship by the Herchel Smith Fund at the University of Cambridge (ET).

## Author Contributions

Conceptualization BW, GK, MF, NR, ET

Investigation BW, AO, MA, SH, JH, SY, ET, LV, MB, EU, JV, LR, PP

Formal analysis BW, AO, MA, SH, JH, SY, ET, LV, MB, EU, JV, LR, PP

Methodology BW, AO, MA, SH, JH, SY, ET, LV, MB, JV, PP, MF

Validation BW, AO, MA, SH, JH, SY, ET, LV, MB, EU, JV, LR, PP

Visualization BW, AO, MA, SH, ET, LV, JV, LR

Data Curation BW, AO, MA, SH, SY, LV, JV, LR

Software AO, JH

Writing – original draft preparation BW, GK,MF, MA, NR

Writing – review and editing BW, GK, MF, BS, MV, NR, BI, AO, MA, SH, JH, SY, ET, LV, EU, JV, LR

Project administration GK, MF, BS, MV, NR, BI

Supervision GK, MF, BS, MV, NR, BI

Resources GK, MF, BS, MV, NR, BI

Funding acquisition GK, MF, BS, MV, NR, BI

## Materials & Correspondence

Further information and requests for resources and reagents should be directed to and will be fulfilled by the Lead Contact, Geert Kops

## Methods

### Data availability

The authors declare that the data supporting the findings of this study are available within the paper and its supplementary information. The ICR1000 UK exome series data are available at the European Genome-phenome archive (EGA), reference number EGAS00001000971. Exome data for individual patients cannot be made publicly available for reasons of patient confidentiality. Qualified researchers may apply for access to these data, pending institutional review board approval.

HeLa cell RNAseq data is available at the NCBI Gene Expression Omnibus database, accession number GSE143392. RNAseq data from the patient and control subject cannot be made publicly available for reasons of patient confidentiality. Qualified researchers may apply for access to these data, pending institutional review board approval.

### Samples

The MVA exome analyses were approved by the London Multicentre Research Ethics Committee (05/MRE02/17). DNA was extracted from whole blood using standard protocols. RNA was extracted from EBV-transformed lymphoblastoid cell lines (LCL) using the RNeasy Mini Kit protocol (QIAGEN).

For the functional experiments the following patient LCLs were used: ID_ 5728_1 (patient, biallelic CENATAC (CCDC84) mutations, ECACC ID: FACT5728DLB), ID_ 5728_3 (sibling, no CENATAC mutations, ECACC ID: FACT5728KC), ID_ 5728_4 (father, monoallelic CENATAC mutation, ECACC ID: FACT5728GLB), ID_ 5728_5 (mother, monoallelic CENATAC mutation, ECACC ID: FACT5728ALB).

LCLs were cultured in RPMI supplemented with 15% fetal bovine serum (FBS), 100 μg/ml penicillin/streptomycin and 2 mM alanyl-glutamine. Cells expressing H2B-mNeon were created by lentiviral transduction, using standard procedures. Imaging of LCLs was performed as previously described^8^.

**Exome sequencing, alignment and variant calling, reference data sets, PTV prioritization method, recessive analysis, & sanger sequencing:** as previously described^8^.

### cDNA analysis of CENATAC (CCDC84) mutations

We synthesised cDNA using the ThermoScript RT-PCR System (Life Technologies) with random hexamers and 1 μg of total RNA. We amplified the mutation regions using cDNA-specific primers and sequenced the PCR products as described above. Primer sequences are available on request.

### Conservation logos

Hidden Markov model (HMM) profiles were created from iterative jackhmmer searches^68^ (version: HMMER3/f [3.1b2 | January 2014]) with CENATAC’s protein sequence against the sequences of all metazoan species within the uniprot database. In-between successive iterations, non-CENATAC sequences were manually removed. Logos were created using Skylign^69^; letter height: information content above background.

### Immunoblots

For the data in Figs. 1C, S4B-C, and S6, cells were treated as indicated and lysed in Laemmli lysis buffer (4% SDS, 120 mM Tris pH 6.8 and 20% glycerol). Lysates were processed for SDS-polyacrylamide gel electrophoresis and transferred to nitrocellulose membranes. Immunoblotting was performed using standard protocols. Visualization of signals was performed on an Amersham Imager 600 scanner using enhanced chemiluminescence. Primary antibodies used were rabbit anti-CENATAC (CCDC84, Sigma, HPA071715) and mouse anti-Tubulin (Sigma, T5168). Secondary antibodies used were goat anti-mouse HRP (170-6516) and goat anti-rabbit HRP (170-6515), both obtained from Bio-Rad.

### HeLa cell culture

HeLa T-REx Flp-In osTIR-9Myc::NEO cells (gift from Andrew Holland) were grown in DMEM high glucose supplemented with 10% Tetapproved FBS, 100 μg/ml penicillin/streptomycin, and 2 mM alanyl-glutamine. Stable expression of H2B-mNeon was done by lentiviral transduction using standard procedures.

### Creation of HeLa EGFP-AID-CENATAC and EGFP-CENATAC cell lines

HeLa EGFP-AID-CENATAC and HeLa EGFP-CENATAC cell lines were derived from HeLa T-REx Flp-In osTIR-9Myc::NEO and HeLa T-REx Flp-In, respectively. Tagging of the endogenous locus of CENATAC was done according to the scCRISPR protocol^70^ using the Protospacer, HDR_insert, and HDR_ext primers in Table S9. pcDNA5-FRT-TO-EGFP-AID (Addgene, 80075) was used as template for both the EGFP-AID and EGFP tags. Cells were transfected with Lipofectamine LTX using standard procedures and subsequently FACS sorted (single cells) based on EGFP expression. Endogenous tagging was confirmed by PCR (using the Genomic primers, Fig. S4A) and immunoblotting of CENATAC protein (Figs. S4B-C).

### Viral plasmids, cloning and virus production

For lentiviral re-expression of CENATAC variants, first pcDNA5 PURO FRT TO EGFP-AID-CENATAC was created by cloning CENATAC cDNA derived from HeLa cells into empty pcDNA5-FRT-TO-EGFP-AID (Addgene, 80075) using the cDNA PCR primers in Table S9 and digestion of both the PCR product and the plasmid with NotI/ApaI. The CENATAC cDNA was subsequently cloned into pcDNA5 PURO FRT TO containing a LAP-tag to create pcDNA5 PURO FRT TO LAP-CENATAC by Gibson assembly^71^ with the PCR primers Gibson1 and Gibson2. Mutagenesis was then performed to make this construct resistant to CENATAC siRNA treatment (CCDC84, Dharmacon, J-027240-07) by Gibson assembly with PCR primers Gibson3. Next, in the siRNA resistant construct, CENATAC wild-type cDNA was mutated to: Mut1 (primers Gibson4), Mut2 (Gibson5), 2xZF (Gibson6; two consecutive rounds of cloning), Δ1 (Gibson7), Δ2 (Gibson8), Δ3 (Gibson9), or Δ4 (Gibson10) by Gibson assembly. Lentiviral CENATAC iresRFP constructs were derived from a lentiviral construct encoding fluorescently tagged histone 2B (H2B) and a puromycin-resistance cassette (pLV-H2B-mNeon-ires-Puro)^72^. First, the fluorescently tagged H2B was substituted by CENATAC derived from pcDNA5 PURO FRT TO LAP-CENATAC (see above) by Gibson assembly with PCR primers Gibson11 and digestion by AscI/NheI. Next the puromycin-resistance cassette was substituted by tagRFP by Gibson assembly with PCR primers Gibson12. Finally, all siRNA resistant variants of CENATAC were cloned from their respective pcDNA5 PURO FRT TO LAP-CENATAC plasmids into pLV CENATAC ires-tagRFP by Gibson assembly with PCR primers Gibson13 and PstI digestion of the plasmid. Virions were generated by transient transfection of HEK 293T cells with the transfer vector and separate plasmids that express Gag-Pol, Rev, Tat and VSV-G. Supernatants were clarified by filtration.

### Immunoprecipitation

For each sample a full 10 cm plate of HeLa EGFP-AID-CENATAC cells was used, treated as indicated (Fig. S4C). The cells were lysed in ice-cold lysis buffer (50 mM Tris pH 7.5, 150 mM NaCl, 2% NP-40, 0.1% deoxycholate, proteasome inhibitors) and treated with benzonase for 15 minutes at 4°C. After centrifugation, the supernatant was incubated with beads (GFP-trap, ChromoTek) for 2.5 hours at 4°C and washed three times with ice-cold lysis buffer. The samples were finally eluted in Laemmli sample buffer.

### Live cell imaging analysis of mitotic fidelity

LCLs were imaged as previously described^8^. HeLa cells were transfected (RNAiMAX, thermofisher) with 40 nM CENATAC siRNA (CCDC84, Dharmacon, J-027240-07) and 1mM 3-indoleacetic acid (IAA) in ethanol or 40 nM GAPDH siRNA (Dharmacon, D-001830-01-05) and ethanol (IAA vehicle) for 24 hours in a 24 well-plate before they were re-plate into 8-well Ibidi μ-slides with 2 mM thymidine (for early S-phase synchronization) and 100 μl virus for CENATAC expression. After 18 hours the cells were released from thymidine for 6 hours and imaged in CO2 independent medium in a heated chamber (37°C), while air-tight sealed in the well-plate. For the experiments in Figs. 2D-E, the cells were incubated with 200 nM SiR-tubulin dye (Spirochrome) for 6 hours prior to imaging to facilitate visualization of the mitotic spindle. Images were acquired every 3 or 5 minutes at 1×1 binning in 7× 2.5 μm z-stacks and projected to a single layer by maximum-intensity projection using NIS-Elements software 4.45. Imaging was performed with a Nikon Ti-Eclipse widefield microscope equipped with an Andor Zyla 4.2 sCMOS camera, 40x oil objective NA 1.3 WD 0.2 mm, and Lumencor SpectraX light engine. Analysis of these experiments was carried out with ImageJ software. When applicable, cells re-expressing CENATAC variants were identified through co-expression of cytosolic RFP (via ires-tagRFP); RFP-negative cells were omitted from the quantifications (see also Fig. S5).

### Immunofluorescence imaging

After treating the cells with siRNAs and IAA (see above) for 24 hours in a 24 well-plate, the cells were re-plated on round 12 mm coverslips and treated with 2 mM thymidine (for early S-phase synchronization) for 24 hours. 10 hours after release MG132 was added for 45 minutes after which the cells were pre-extracted with 0.1% Triton X-100 in PEM (100 mM PIPES pH 6.8, 1 mM MgCl2 and 5 mM EGTA) for ±60 seconds. After 60 seconds 4% paraformaldehyde was added on top of the PEM in a 1:1 ratio (400 μl each) for 20 minutes to fixate the cells. The coverslips were subsequently washed twice with PBS and blocked with 3% BSA in PBS for 16 hours at 4°C, incubated with primary antibodies for 2 hours at room temperature, washed 3 times with PBS containing 0.1% Triton X-100, and incubated with secondary antibodies for 1 hour at room temperature. Coverslips were then washed 4 times with PBS/0.1% Triton X-100 and mounted using ProLong Gold Antifade with DAPI (Molecular Probes). All images were acquired on a deconvolution system (DeltaVision Elite; Applied Precision/GE Healthcare) equipped with a 100x/1.40 NA UPlanSApo objective (Olympus) using SoftWorx 6.0 software (Applied Precision/GE Healthcare). The images are maximum intensity projections of deconvoluted stacks. Random pro-metaphase and metaphase cells were selected and centrioles were counted by hand. Primary antibodies used were rabbit anti-Centrin1 (Abcam, ab101332, 1/500) and mouse anti-Tubulin (Sigma, T5168, 1/10.000). Secondary antibodies used were goat anti-mouse 647 (A21236) and goat anti-rabbit 568 (A11036), both obtained from Thermofisher.

### Co-evolution analysis

First, a phylogenetically diverse set of complete eukaryotic predicted proteomes was utilized. This set was previously compiled to contain the protein sequences of 90 eukaryotic species^43,73^. These species were selected based on their representation of eukaryotic diversity. If available, we selected two species per clade and model organisms were preferred over other species. If multiple proteomes or proteomes of different strains were available, the most complete proteome was selected. When multiple splicing variants of a single gene were annotated, the longest protein was chosen. A unique protein identifier was assigned to each protein, consisting of four letters and six numbers. The letters combine the first letter of the genus name with the first three letters of the species name. The versions and sources of the selected proteomes can be found in Table S1.

To define phylogenetic profiles for all human proteins, we determined automatic orthologous groups (OG) across the database using information from PANTHER 9.0^74^. PANTHER 9.0 contains 85 genomes with in total 1136213 genes. Of these genes, 759627 genes are in PANTHER families with phylogenetic trees, multiple sequence alignments and HMM profiles. In total there are 7180 PANTHER families and 52768 subfamilies. Families are groups of evolutionary related proteins and subfamilies are related proteins that are likely to have the same function. The division into subfamilies is done manually, by biological experts. Every subfamily of PANTHER is an OG at some taxonomic level in the tree of life. We used 'hmmscan' tool from the HMMER package^68^ (HMMER 3.1b1) to find for each protein sequence in our database, the best matching profile of a mainfamily or subfamily in PANTHER9.0. The phylogenetic profile of panther main- or subfamily was subsequently defined by utilizing the hierarchical nature of the panther classification. Specifically the phylogenetic profile of a main- or subfamily also include all members of daughter families (and if relevant their daughter families etc.). Note that due to the automatic nature of orthology definition and the draft quality of a few genomes, phylogenetic profiles of the human proteins are not as accurate as those defined by manual analysis^75^.

To determine the phylogenetic profile similarity, pearson correlation (https://en.wikipedia.org/wiki/Phi_coefficient) was computed between the phylogenetic profile of the CENATAC panther (PTHR31198) and the phylogenetic profile of all other panther sub and main families using in house scripts. To detect functional patterns in orthologous groups with similar phylogenetic profiles (correlation > 0.5), a GO Enrichment Analysis was performed^76–78^. GO cellular component overrepresentation (GO Ontology database: released 2020-01-03) was computed using PANTHER (test release 2019-07-11) with the human reference genome gene set as background. Statistical significance of overrepresented GO terms was computed using Fisher’s exact test with FDR correction.

### Nuclear extract and GFP pull down and mass spectrometry

Nuclear extract of wild-type and EGFP-CENATAC HeLa cells was prepared as described earlier^79^. In short, cells were harvested by trypsinization and resuspended in cold hypotonic buffer (10 mM Hepes KOH pH 7.9, 1.5 mM MgCl2, 10 mM KCl). Afterwards, the cell pellet was homogenized using a Douncer with type B pestle (tight) to lyse the cell membrane. After centrifuging, the nuclei were washed with cold PBS and resuspended in cold buffer for lysis (420 mM NaCl, 20 mM Hepes KOH pH 7.9, 20% v/v glycerol, 2 mM MgCl2, 0.2 mM EDTA) followed by rotation, centrifugation, and collection of the nuclear extract. 450 μl of nuclear extract was used for each GFP pull down using 15 μl slurry of GFP-Trap agarose beads (Chromotek), performed in triplicate. GFP pull-downs were done as described earlier^80^, without the addition of EtBr during the incubation, and with an adapted buffer C (150 mM NaCl, 20 mM Hepes KOH pH 7.9, 20 % v/v glycerol, 2 mM MgCl2, 0.2 mM EDTA, complete protease inhibitors w/o EDTA, 0.5 mM DTT) for the incubation (+0.1% NP40) and washes (+0.5% NP40). Samples were digested using on-bead digestion with trypsin overnight^81^. The tryptic peptides were acidified with TFA and purified on C18 StageTips^82^.

After elution from the C18 StageTips, tryptic peptides were separated on an Easy-nLC 1000 (Thermo Scientific), connected online to a Q-Exactive HF-X Hybrid Quadrupole-Orbitrap Mass Spectrometer (Thermo Scientific), using an acetonitrile gradient of 7-30% for 48 min followed by washes of 50-90% acetonitrile, for 60 min of total data collection. Full scans were measured with a resolution of 120.000, the top twenty most intense precursor ions were selected for fragmentation with a resolution of 15.000 and dynamic exclusion set at 30 sec. Peptides were searched against the UniProt human proteome (downloaded June 2017) using MaxQuant^83^ (version 1.6.0.1) with default settings, and iBAQ, LFQ, and match-between-runs enabled. Data analysis was done using Perseus (version 1.5.5.3), the volcano plot and stoichiometry calculations were done as described earlier^80^ using in-house made scripts for R (version 3.6.1).

### Nuclear Extract Preparations for Northern blots and glycerol gradients

Nuclear extract from Hela cells were prepared according to the protocol described by Dignam et al^84^ using buffer D containing 50 mM KCL in the final dialysis step.

### Immunoprecipitation and Northern blots

100 μl nuclear extract diluted in lysis buffer to a final volume of 200 μl was incubated with 2μg of anti-CCDC84 antibody (SIGMA-HPA071715) overnight in the cold room with end-to-end rotation. The following day capture of antibody-antigen complexes was done using 50 μl of resuspended Protein G Dynabeads prepared according to manufacturer’s instructions and incubated with the nuclear extract-antibody samples for 2 hr at 4°C. Beads were then washed four times with lysis buffer lacking protease and RNase inhibitors. RNA was eluted by proteinase K treatment, extracted once with phenol:chloroform:isoamyl alcohol (25:24:1; pH4.8) followed by ethanol precipitation. RNA dissolved to H2O or 0.1X TE buffer.

Total volumes of 2 μl (Input) and 5 μl (IP) RNA samples were separated on a 6% polyacrylamide-Urea gel and analysed by Northern blotting essentially as described in^55^. Individual snRNAs were detected using 32P 5’-end labeled DNA or LNA oligonucleotides complementary to individual snRNAs. Northern blots were exposed to image plates and visualized using Typhoon FLA-9400 scanner (GE Healthcare, US) at 50 micron resolution. The data was quantified using AIDA software (Raytest, Germany).

### Glycerol Gradient and ultracentrifugation

Nuclear extracts were preincubated for 0-20 min at +30 °C in a buffer containing 13 mM HEPES(pH 7.9), 2.4 mM MgCl2, 20 mM creatine phosphate, 2 mM DTT, 40 mM KCl, 0.5 mM ATP. Aggregates were subsequently removed by a brief centrifugation (20 000 g, 1 min, +4°C) and the supernatant subsequently ultracentrifuged on a linear 10-30% gradient (20 mM HEPES, pH 7.9; 40 mM KCl, 2 mM DTT, 2.4 mM MgCl2) for 18 hr at 29000 rpm, +4°C, Sorvall TH641 rotor. Following ultracentrifugation, the samples were fractionated. 20% of each fraction was deproteinized and used for RNA isolation and Northern blotting and the remaining 80% was subjected to TCA precipitation, separated on a 10% SDS-PAGE and analysed by Western blots. Each blot was probed for CENATAC (CCDC84-HPA071715, Sigma-Aldrich-Merck), PRPF4 (#HPA0221794, Sigma-Aldrich-Merck) and TXNL4B (E-AB-61535, Elabscience, US).

### RT-PCRs

For Fig. 4A: total cellular RNA was extracted using the RNeasy kit protocol (Qiagen) and treated with DNase I Amplification grade (Invitrogen) to remove potential genomic DNA contamination. cDNA synthesis was carried out using SuperScript™ II RT (Thermo Fisher Scientific) and Oligo(dT)18 primers. PCRs were performed with Phusion High-Fidelity DNA polymerase (Thermo Fisher Scientific) with the following cycling conditions: initial denaturation (98°C for 60 sec), followed by either 30 cycles of denaturing (98°C for 10 sec), annealing (gene-specific temp for 30 sec), extension (72°C for 30 sec) and a final extension (72°C for 1 min 30 sec). PCR primers and relevant annealing temperatures are listed in Table S9. PCR products were analyzed on 2% agarose gel run using 1X TBE buffer. For Figs. 4B and 5E total RNA isolated was isolated from HeLa cells or patient/control subject lymphoblasts using Trizol extraction followed by an additional acidic phenol (pH 5.0) extraction. 1 μg of RNA was converted to cDNA using Maxima H minus reverse transcriptase (Thermo Fisher) according the manufacturer protocol. PCRs were performed essentially as described above and gene specific primers and (annealing temperatures) are listed in Table S9.

### RNA isolation and high-throughput sequencing

Total RNA isolated was isolated from EGFP-AID-CENATAC HeLa cells treated with siGAPDH (Dharmacon, D-001830-01-05) for 48 hours or with siCENATAC (CCDC84, Dharmacon, J-027240-07) and 1mM 3-indoleacetic acid (IAA) for 24 or 48 hours, or unedited HeLa parental cells treated with siGAPDH for 48 hours, or patient/control subject lymphoblasts using Trizol extraction followed by an additional acidic phenol (pH 5.0) extraction. RNAseq libraries were constructed using Illuminan TruSeq Stranded Total RNA kit (Illumina) Human Ribo-Zero rRNA depletion kit (Illumina). Paired-end 150+150 bp sequencing was done with Illumina NextSeq 500 using NextSeq 500/550 High Output Kit v2.5 for HeLa samples and with Illumina NovaSeq 6000 using partial S4 flow cell lane for patient samples.

### Mapping the reads to the genome

The STAR aligner^85^ was used for mapping the paired sequence reads to the genome (hg38/GRCh38). Transcript annotations were obtained from GENCODE (v29). The length of genomic sequence flanking the annotated junctions (sjdbOverhang parameter) was set to 161. The Illumina adapter sequences AGATCGGAAGAGCACACGTCTGAACTCCAGTCAC and AGATCGGAAGAGCGTCGTGTAGGGAAAGAGTGTAGATCTCGGTGGTCGCCGTATCATT were, respectively, clipped from the 3’ of the first and the second pairs in the read libraries (using clip3pAdapterSeq parameter).

### Differential alternative splicing analysis

Differential alternative splicing (AS) analysis was done using Whippet (v0.11)^51^. Both merged aligned reads (bam files) and AS event annotations from GENCODE (v29) were used to build the index reference for AS events. To detect the significantly differential events, probability cutoff of Pr > 0.9 and Percentage Spliced In deviation cutoff of |ΔΨ| > 0.1 were used.

### Differential intron retention analysis

For a comprehensive and sensitive intron retention (IR) analysis the IntEREst R/Bioconductor package was used^50^. After reading binary alignment (.bam) files, IntEREst detects introns with significantly higher and lower number of mapped reads relative to the number of reads that span the introns. The DESeq2-based function of IntEREst, i.e. deseqInterest(), was used for the differential IR analysis. The Benjamini-Hochberg method was used for adjusting the p-values and a cutoff of padj<0.05 was applied to extract the significantly differential IRs. The reference table was built from the NCBI RefSeq transcription annotations based on hg38/GRCh38 genome assembly.

### Annotating U12-type introns

We used IntEREst R/Bioconductor package to annotate the U12-type introns as described previously^50^ using threshold values of 0.07 and 0.14 for 5’ss and BPS scores, respectively. BPS sequence was identified by scanning intronic region from position −40 to position −3 upstream of the 3’ss and the highest scoring sequence was selected as the BPS. This list was manually appended with additional introns that did not fulfil our annotation criteria (typically because of poor BPS sequence), but have been previously identified as U12-type introns^86^.

### P120 minigene cloning, transfection and analysis of RNA

The double 5’ss constructs were created by insertion mutagenesis PCR using the P120 minigene^53^ as a template, and further modifications of 5’ splice sites were made by PCR using mutagenic primers (for a list of primers used see Table S9). The 3’ss was modified to accommodate for GT-subtype splicing by insertion of a CAG trinucleotide sequence through insertion mutagenesis PCR. All mutations were confirmed by DNA sequencing. Chinese Hamster Ovary cells were transfected with the double 5’ss constructs (1600 ng per well of a 12-well plate) using Lipofectamine 2000 (ThermoFisher Scientific) and after 24h total RNA was isolated using TRIZOL reagent (ThermoFisher Scientific). Following DNAse treatment, a pCB6 vector specific oligonucleotide (ACAGGGATGCCA) was used for reverse transcription of the RNA with Revertaid (ThermoFisher Scientific). RT-PCR was performed with primers binding exon 6 (GGATGAGGAACCATTTGTGC) and exon 7 (AGAACGAGACCGCCCTTC), and the resulting PCR products were analyzed on a 3 % MetaPhorTM (Lonza) agarose gel. The gel was imaged using Fuji LAS-3000 CCD camera and the band intensities were quantified using AIDA software (Raytest, Germany). Identities of the PCR products were confirmed by DNA sequencing.

## Supplemental Information

1. **Supplementary Figures**
2. **Movie S1.** GAPDH depletion EGFP-AID-CENATAC HeLa cells expressing H2B-mNeon (upper) and depleted of GAPDH. Microtubules were visualized with SiR-Tubulin (lower). Time in hours.
3. **Movie S2.** CENATAC depletion EGFP-AID-CENATAC HeLa cells expressing H2B-mNeon (upper) and depleted of CENATAC. Microtubules were visualized with SiR-Tubulin (lower). Time in hours.
4. **Table S1.** List of species used in the co-evolution analysis of Fig. 3A with additional information and references.
5. **Table S2.** Co-evolution scores List of co-evolution correlation scores per Panther/Ensembl Gene ID.
6. **Table S3.** CENATAC depletion intron retention dataset CENATAC RNAseq data analyzed for intron retention using IntEREst and for alternative splicing using Whippet. Sequencing statistics and data for the subtypes of U2- and U12-type introns is shown in separate tabs.
7. **Table S4.** MDS intron retention dataset MDS RNAseq data (GEO GSE63816) analyzed for intron retention using IntEREst and for alternative splicing using Whippet. Sequencing statistics and data for the subtypes of U12-type introns is shown in separate tabs.
8. **Table S5.** MVA patient intron retention dataset CENATAC RNAseq data analyzed for intron retention using IntEREst. Sequencing statistics and data for the subtypes of U2- and U12-type introns is shown in separate tables.
9. **Table S6.** Venn diagram data related to Figure 5G.
10. **Table S7.** Go-term enrichment statistics related to Figure S15.
11. **Table S8.** Full metascape results table related to Figure S15.
12. **Table S9.** List of primers used.

---

**Fig. S1.**
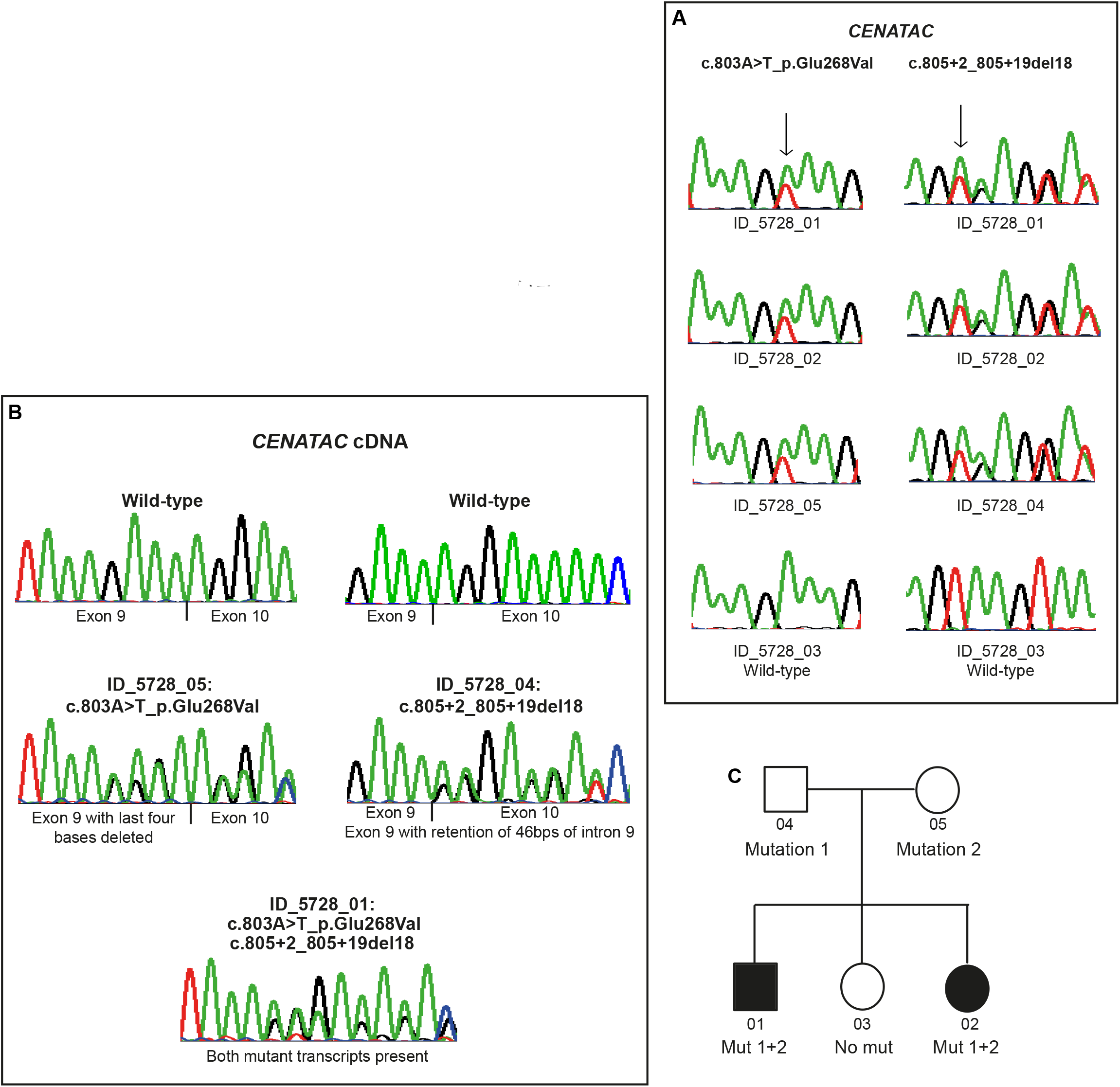
Case report and chromatograms of individuals with mutations in *CENATAC* (*CCDC84*). **A)** Sequencing chromatograms showing mutations in blood DNA and corresponding wild-type sequence from a control. **B)** Sequencing chromatograms from Reverse Transcription-PCR analysis of RNA showing the effect of *CENATAC* mutations. Maternal cDNA sequencing (ID_5728_05) demonstrates that c.803A>T_p.Glu268Val leads to a translational frameshift as a result of deletion of the last four bases of exon 9. Paternal cDNA sequencing (ID_5728_04) shows that c.805+2_805+19del18 results in retention of 46 bps of intron 9. The affected child’s cDNA sequencing (ID_5728_01) demonstrates both mutant transcripts are present. **C)** Pedigree of family (ID_5728) showing CENATAC mutation status.

**Fig. S2.**
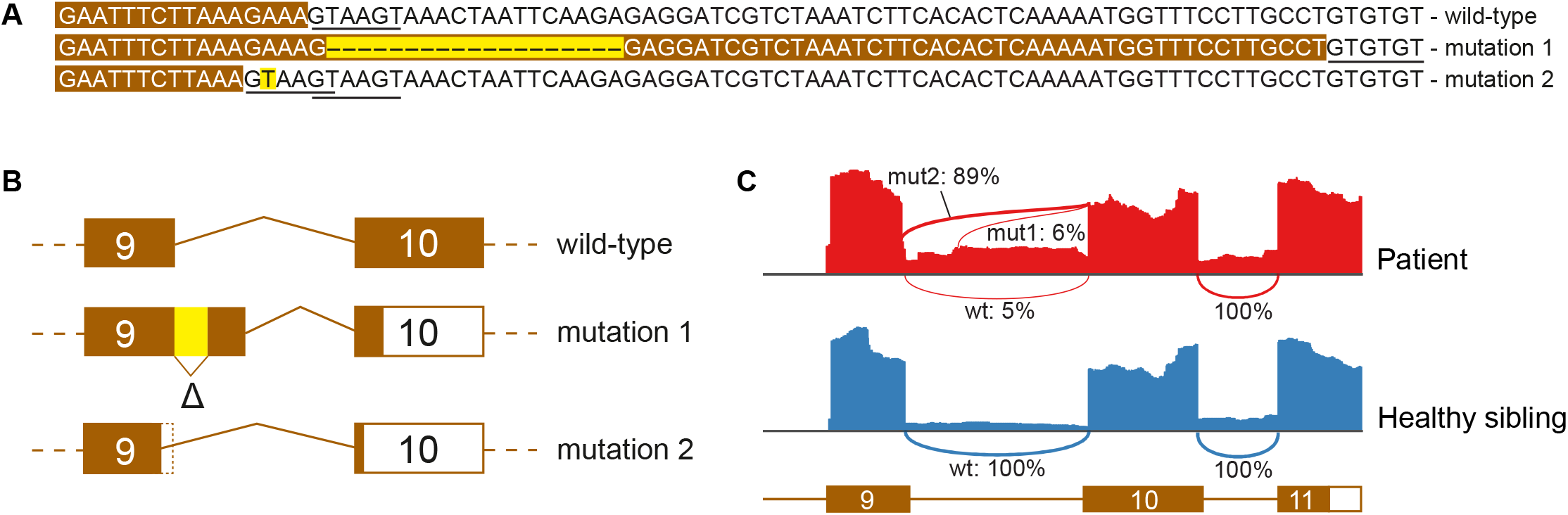
The effect of the MVA patient mutations on splicing of *CENATAC* intron 9. **A)** DNA sequences of exon 9 (white in brown boxes) and intron 9 (black) of wild-type and MVA mutated *CENATAC* alleles. The mutations (c.805+2_805+19del18, mut1; and c.803A>T, mut2) are indicated with yellow. Both the new and original 5’ splice sites are underlined. **B)** Schematic showing the effect of the MVA patient mutations on splicing of CENATAC intron 9. Exons 9 and 10 are represented with rectangles; protein-coding regions within exons are indicated with fill, untranslated regions without fill. See also Fig. 1C. **C)** Sashimi plots derived from patient and healthy sibling RNAseq data showing showing the effect of the MVA patient mutations on 5’ splice site useage (percentages) of intron 9 in patient cells. The different splice sites are shown in A).

**Fig. S3.**
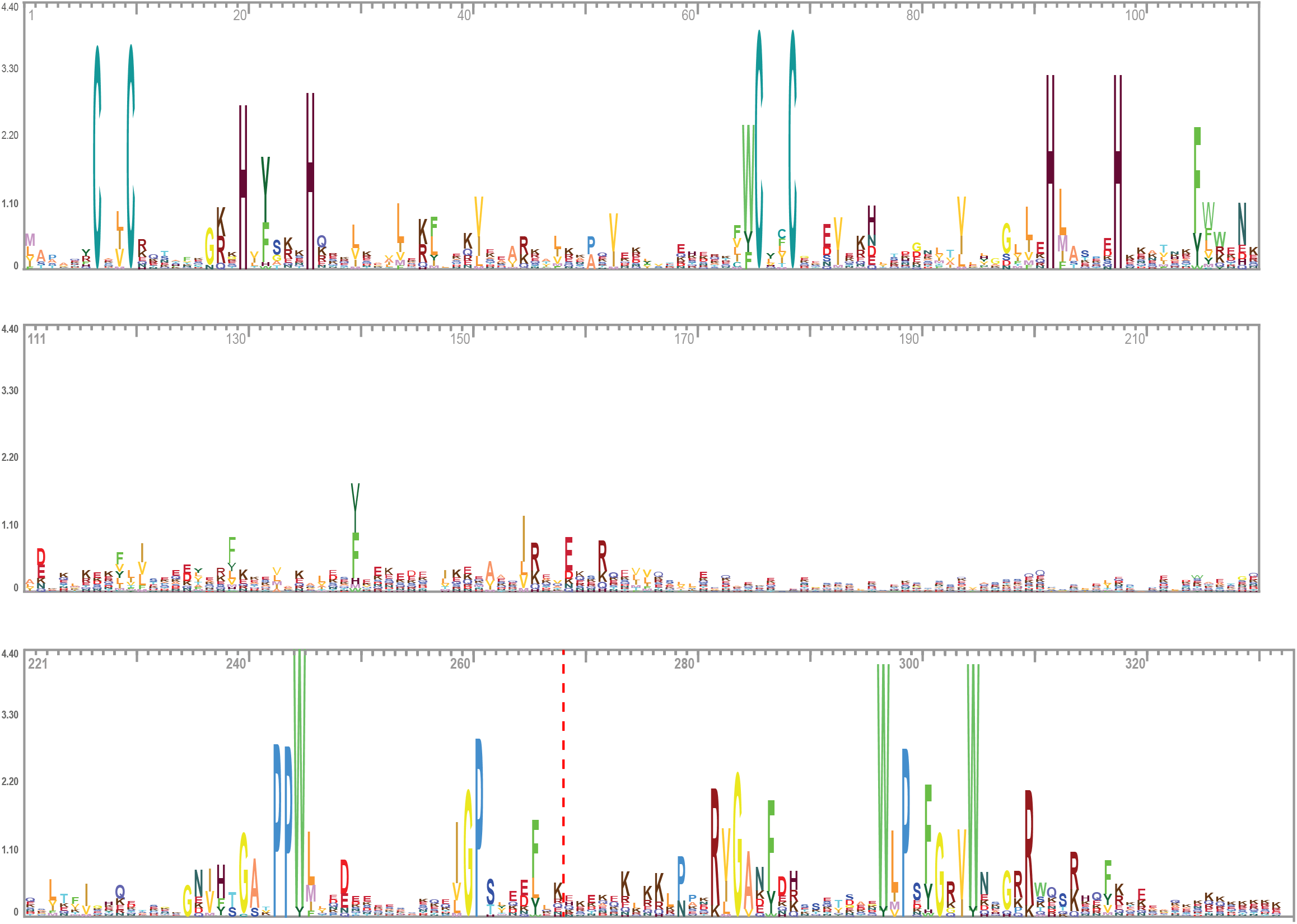
Full-length sequence logo of CENATAC’s conserved residues (in metazoan species) split in three. Amino acids are numbered. The location of the truncating MVA mutations is indicated with the red dotted line.

**Fig. S4.**
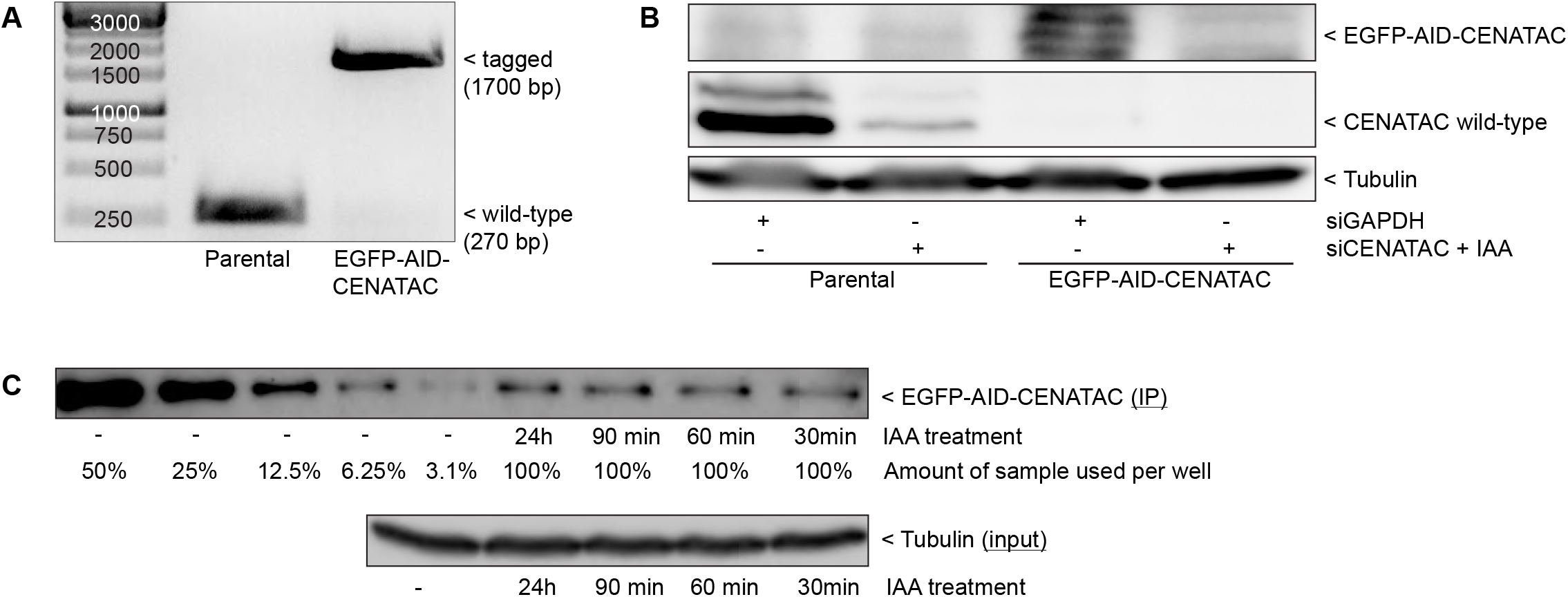
Endogenous EGFP-AID-CENATAC tag verification and CENATAC depletion. **A)** Genomic PCR of the CENATAC locus around the start codon. Left: parental cells; right: EGFP-AID-CENATAC cells. **B)** CENATAC (αCENATAC) and tubulin (αTubulin) immunoblots of EGFP-AID-CENATAC and parental cells treated as indicated for 48 hours. **C)** CENATAC (αCENATAC) immunoblot of EGFP-AID-CENATAC immunoprecipitated from cells treated with IAA for the indicated amounts of time (upper) and Tubulin (αTubulin) immunoblot of input samples (lower). The untreated IP sample (upper gel, 5 left-most wells) was divided into volumes of 50, 25, 12.5, 6.25 and 3.1% of the totoal volume, as indicated, for comparison with the IAA treated samples (4 right-most wells) of which 100% of the total volume was loaded.

**Fig. S5.**
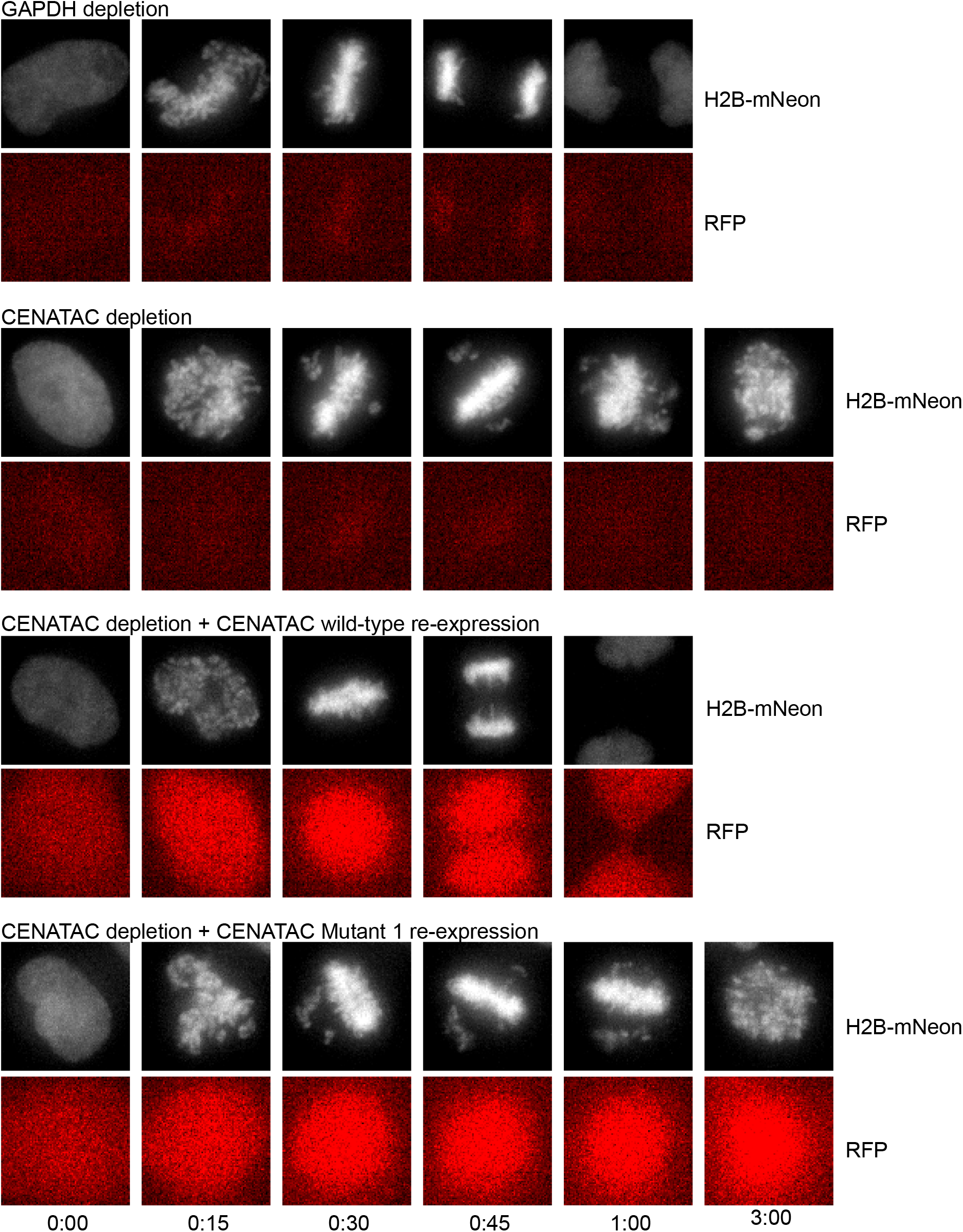
EGFP-AID-CENATAC cells expressing H2B-mNeon depleted of GAPDH or CENATAC, with or without re-expression of CENATAC variants as indicated (as in Figs.1F and 1G). Expression of cytosolic RFP was used as a marker to identify cells expressing CENATAC variants. Upper panels: H2B-mNeon; lower panels: RFP. Time in hours.

**Fig. S6.**
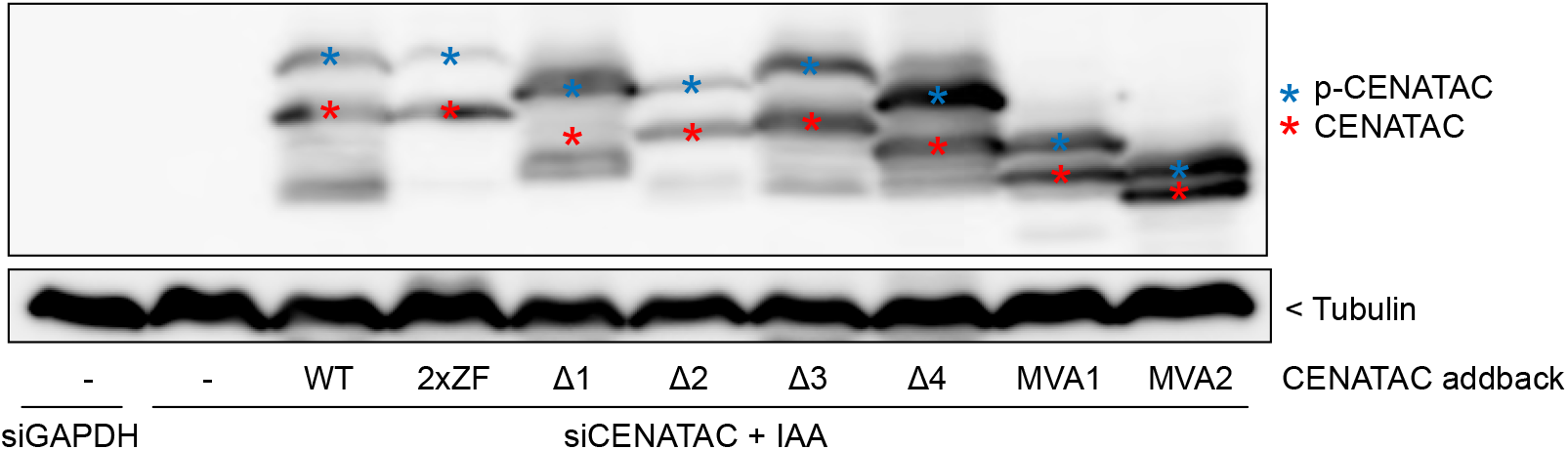
CENATAC (αCENATAC) and tubulin (αTubulin) immunoblots of cells treated as in Fig. 1G. p-CENATAC indicates phosphorylated CENATAC (blue asterisks).

**Fig. S7.**
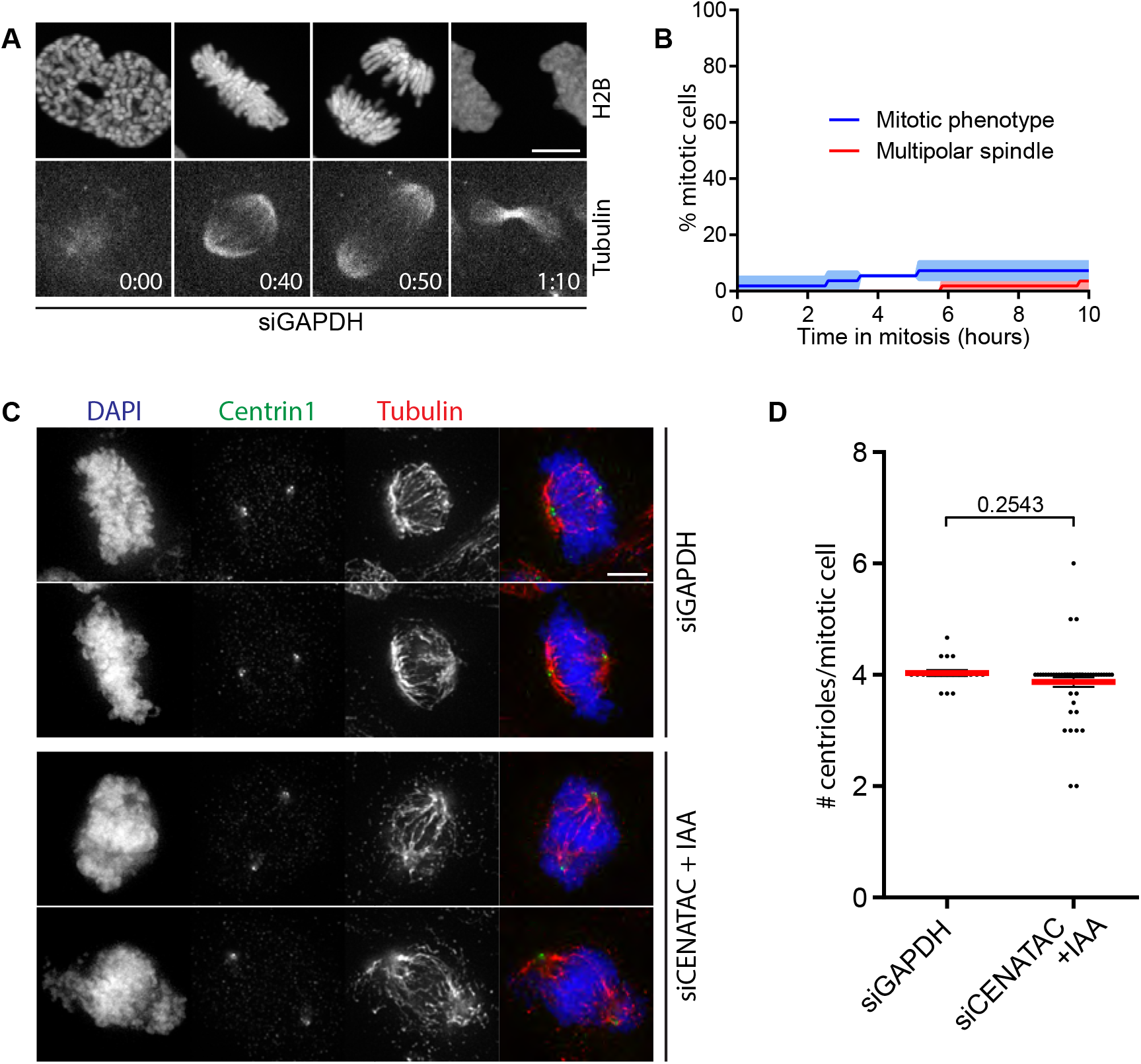
CENATAC’s congression phenotype is not the result of a multipolar mitotic spindle. **A)** Representative stills of EGFP-AID-CENATAC HeLa cells expressing H2B-mNeon and depleted of GAPDH. Microtubules were visualized with SiR-Tubulin. Scale bar, 5 μm. Time in hours. See also Movies S1 and S2. **B)** Quantification of the mitotic phenotype and multipolar spindle formation in time in cells treated as in A). Each line depicts the mean of three experiments ± s.e.m., with >44 cells in total. **C)** Representative immunofluorescence images of EGFP-AID-CENATAC cells depleted of GAPDH or CENATAC and stained with antibodies against Centrin1 and Tubulin. Scale bar, 5 μm. **D)** Quantification of the amount of centrioles per mitotic cell treated as in C). Each line depicts the average of three experiments ± s.e.m., with >60 cells in total. The P value was calculated with a two-sided unpaired Student’s t test.

**Fig. S8.**
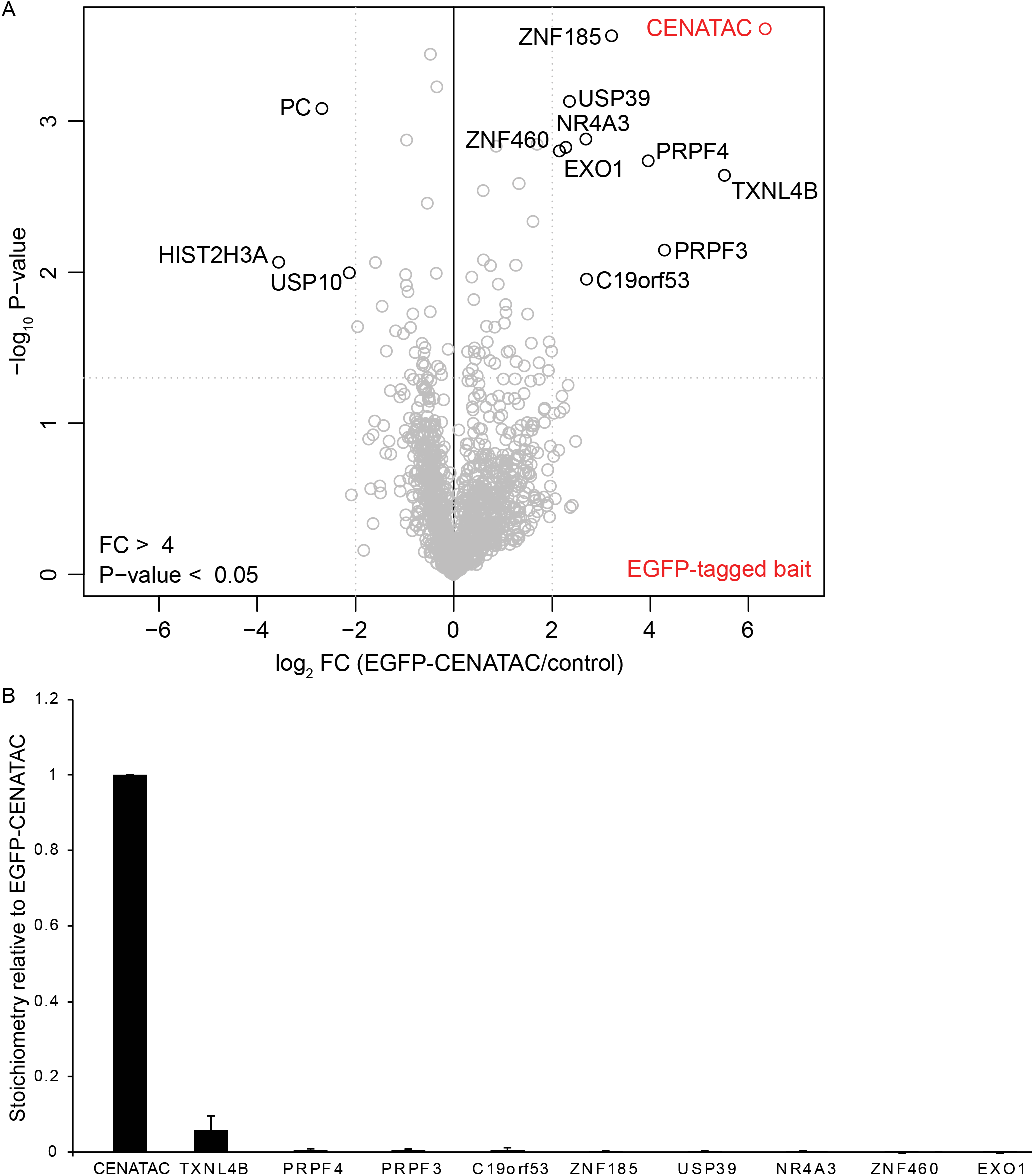
Volcano plot and stoichiometry of EGFP-CENATAC immunoprecipitations. **A)** Mass spec-based identification of interacting proteins with EGFP-CENATAC in HeLa cells. Statistically enriched proteins in the EGFP-CENTAC pull downs (n = 3), determined using a two tailed t-Test, are depicted on the right-hand side of the volcano plot. The fold change of the LFQ intensities, from the CENTAC over the wt pulldown, is depicted on the x-axis (log2). The y-axis shows the −log10 P-value. Cutoffs are shown with a dotted grey line, with al fold change of 4 and a P-value of 0.05. **B)** Stoichiometry of proteins interacting with EGFP-CENATAC. The iBAQ value of each proteinis divided by the iBAQ value of CENATAC, and depicted with CENATAC set to 1. Data are shown as mean ± s.d. (n = 3 pulldowns).

**Fig. S9.**
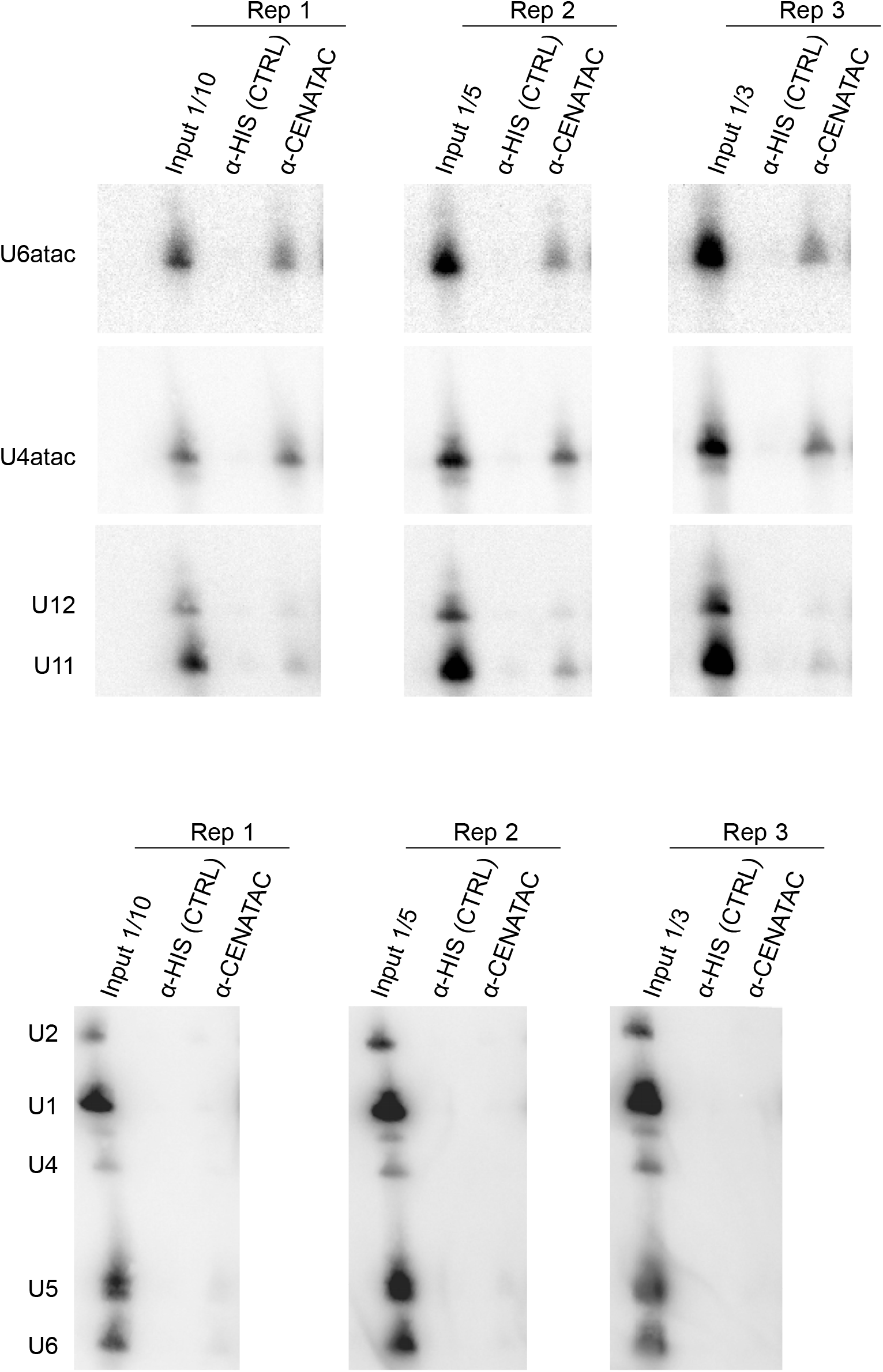
Northern blots of EGFP-CENATAC immunoprecipitations (N=3) used for the quantification in Fig. 3E.

**Fig. S10.**
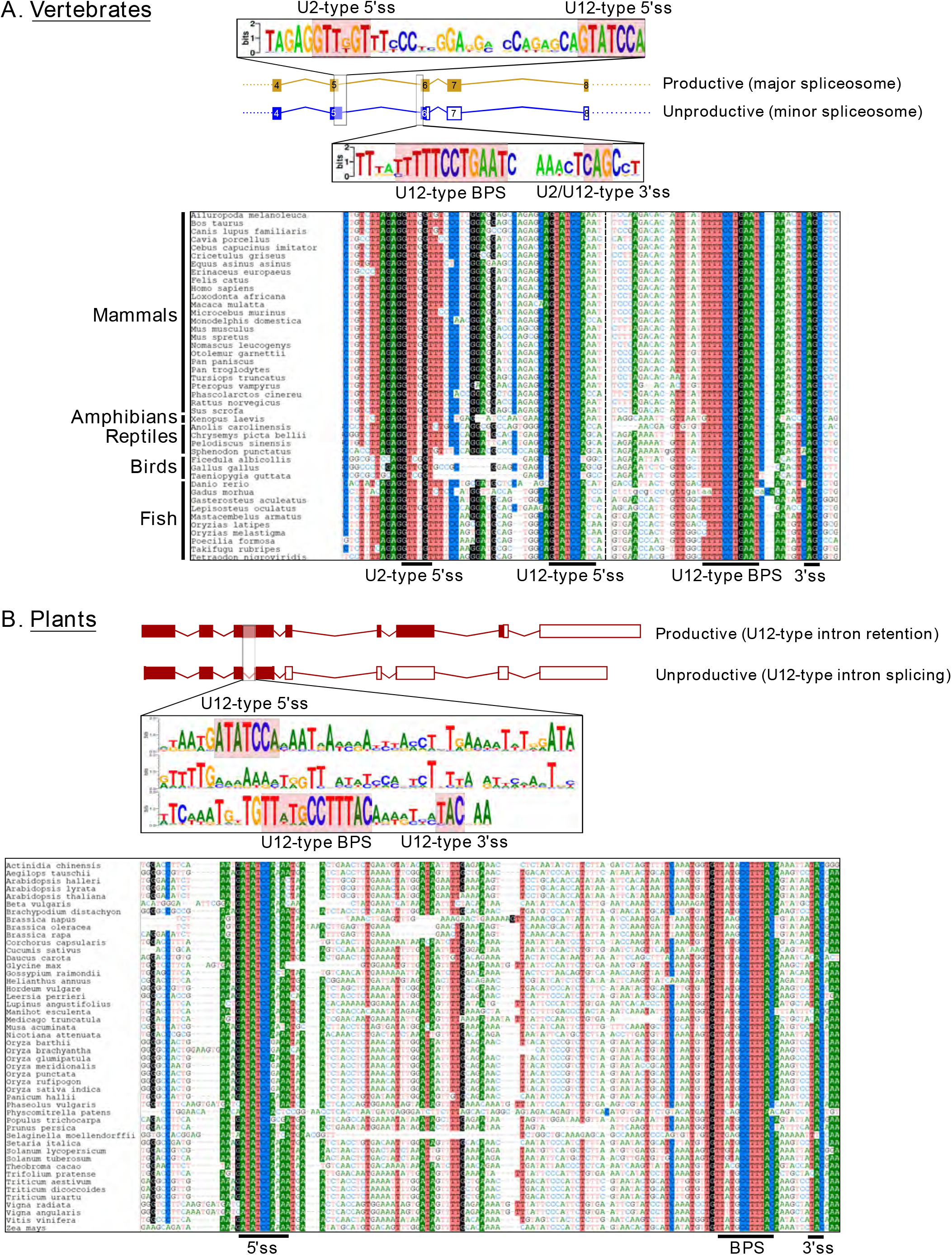
Evolutionarily conserved sequence elements involved in production of CENATAC productive and nonproductive mRNA isoforms in vertebrates and plants. **A)** In vertebrates splicing of intron 5 by the major spliceosome leads to productive mRNA formation which includes the full-length CENATAC coding sequence. Splicing by the minor spliceosome, however, leads to nonproductive mRNA formation due to introduction of a premature termination codon in exon 6. Sequence logos in the upper panel show the conservation of the U2- and U12-type 5’-splice sites, U12-type BPS and the 3’ss (used by both spliceosomes). The schematic shows exon 4-8 of the human transcripts ENST00000334418.6 (productive isoform) and ENST00000532132.5 (nonproductive isoform). The aligned sequences (lower panel) include 43 vertebrate species (25 mammals, 1 amphibian, 3 reptiles, 1 lizard, 3 birds, 10 fish) that were aligned with DiAlign (dialign.gobics.de) followed by a subsequent sequence logo construction using WebLogo (weblogo.berkeley.edu). **B)** In plants retention of a U12-type intron embedded in exon 3 leads to productive mRNA formation that includes the full-length CENATAC coding sequence. Splicing of the U12-type intron by the minor spliceosome leads to a nonproductive mRNA formation due to introduction of premature termination codon in exon 4. The sequence logo shows the conservation of the U12-type 5’ss, BPS and 3’ss elements within the retained intron. In both panels **A)** and **B)** the filled rectangles indicate protein-coding exons, open rectangles represent non-coding exons. Nucleotides showing at least 85% identity between the species are shaded.

**Fig. S11.**
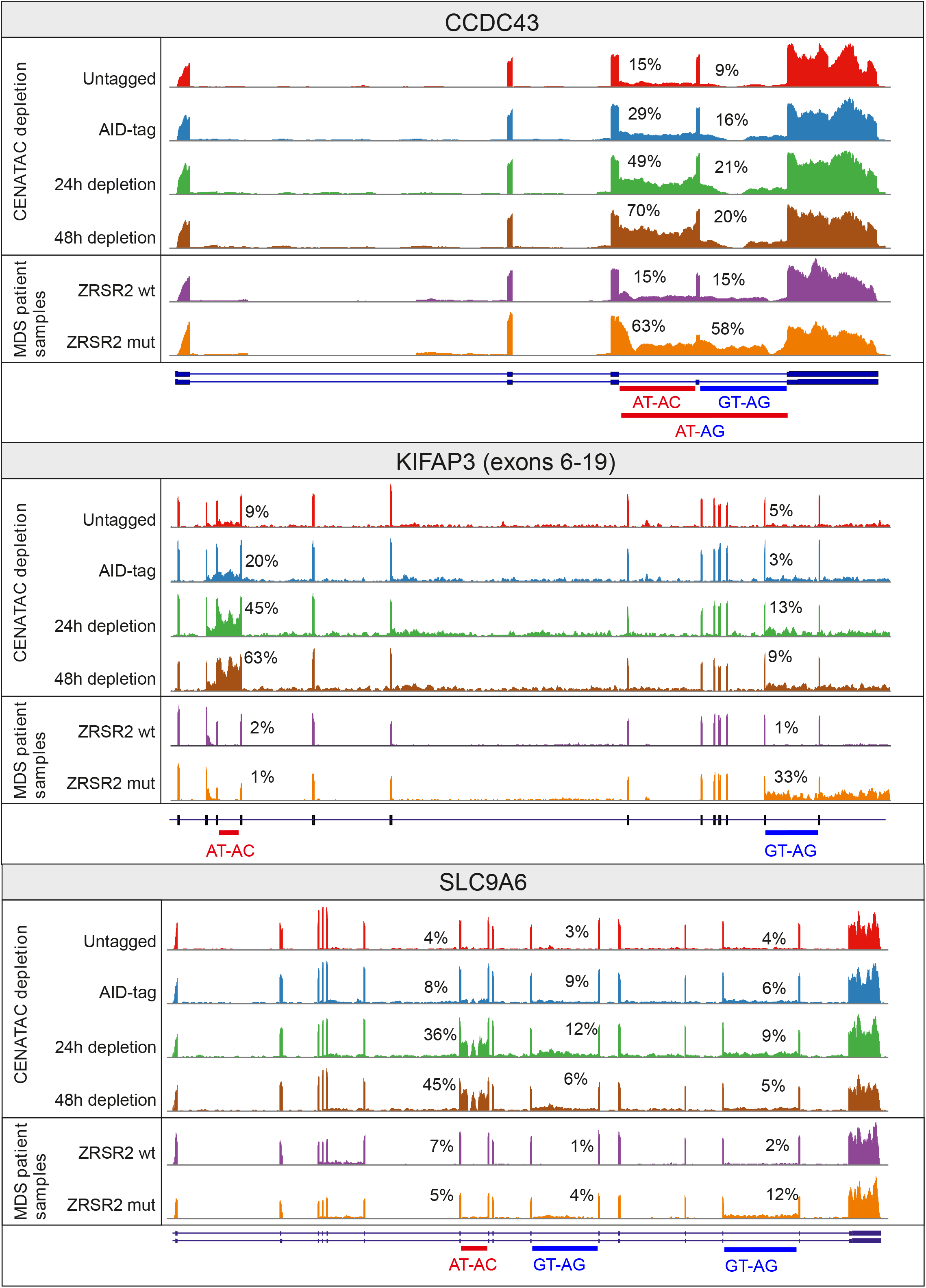
Sashimi plots showing the effect of CENATAC depletion (24 or 48 hours) in EGFP-AID-CENATAC HeLa cells or *ZRSR2* mutations in the MDS dataset on A- and G-type minor intron retention: AT-AC and GT-AG introns, respectively. Untagged represents the parental unedited HeLa cell line, AID-tag represents the EGFP-AID-CENATAC HeLa cell line; both were depleted of GAPDH for 48h. The AT-AG intron (*CCDC43,* in red and blue) is a hybrid intron that starts from 5' splice site of the AT-AC intron and ends with the 3’ splice site of the GT-AG intro.

**Fig. S12.**
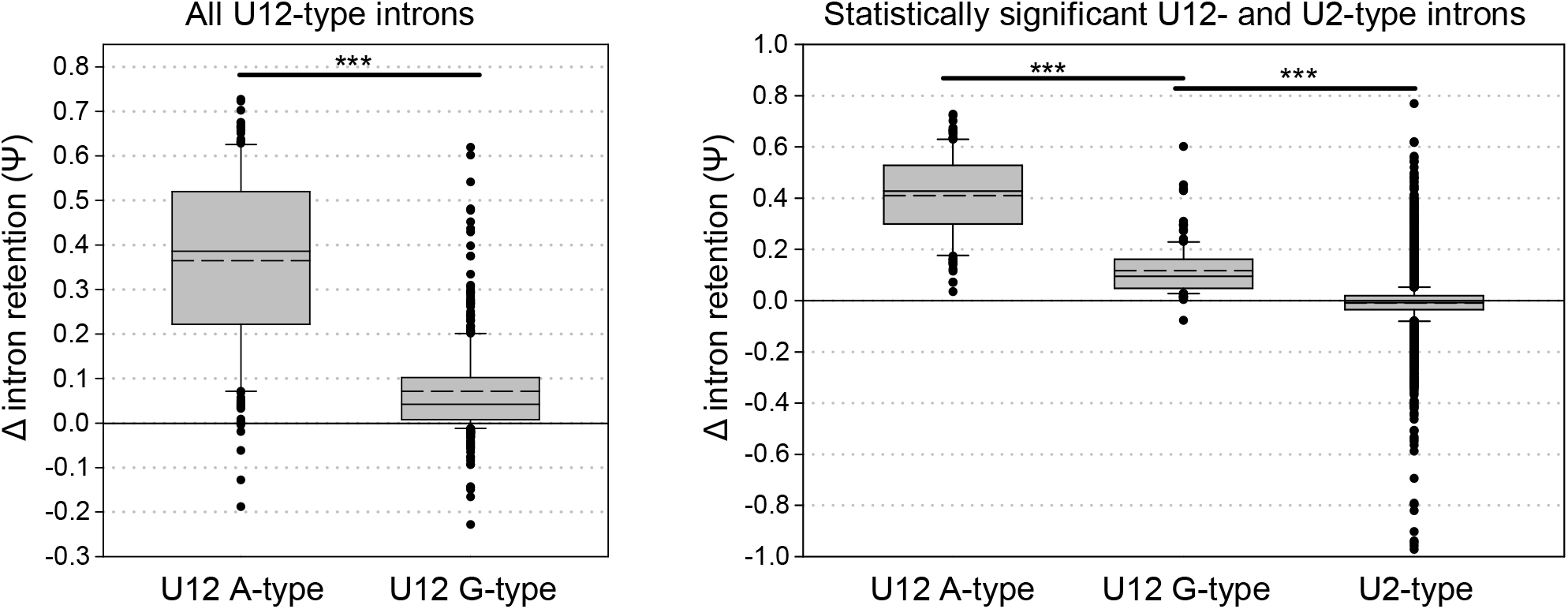
Comparison of the delta-psi values (EGFP-AID-CENATAC HeLa cells 48h CENATAC depletion vs. parental cell line 48h GAPDH depletion). **A)** Comparison of all (significant and not significant) U12 A-type (n=179) and U12 G-type (n=441) introns. **B)** comparison of statistically significant U12 A-type introns (n= 133), U12 G-type introns (n=130), and U2-type introns (n=8818). Only introns with on average at least 5 intron-mapping reads were used in the analysis. The boundaries of the boxes indicate 25th and 75th percentiles. Whiskers indicate the 90th and 10th percentiles. Median is indicated with solid line, mean with dashed line inside the box. *** - P<0.001, ns - P>0.05 (Mann-Whitney Rank Sum Test).

**Fig. S13.**
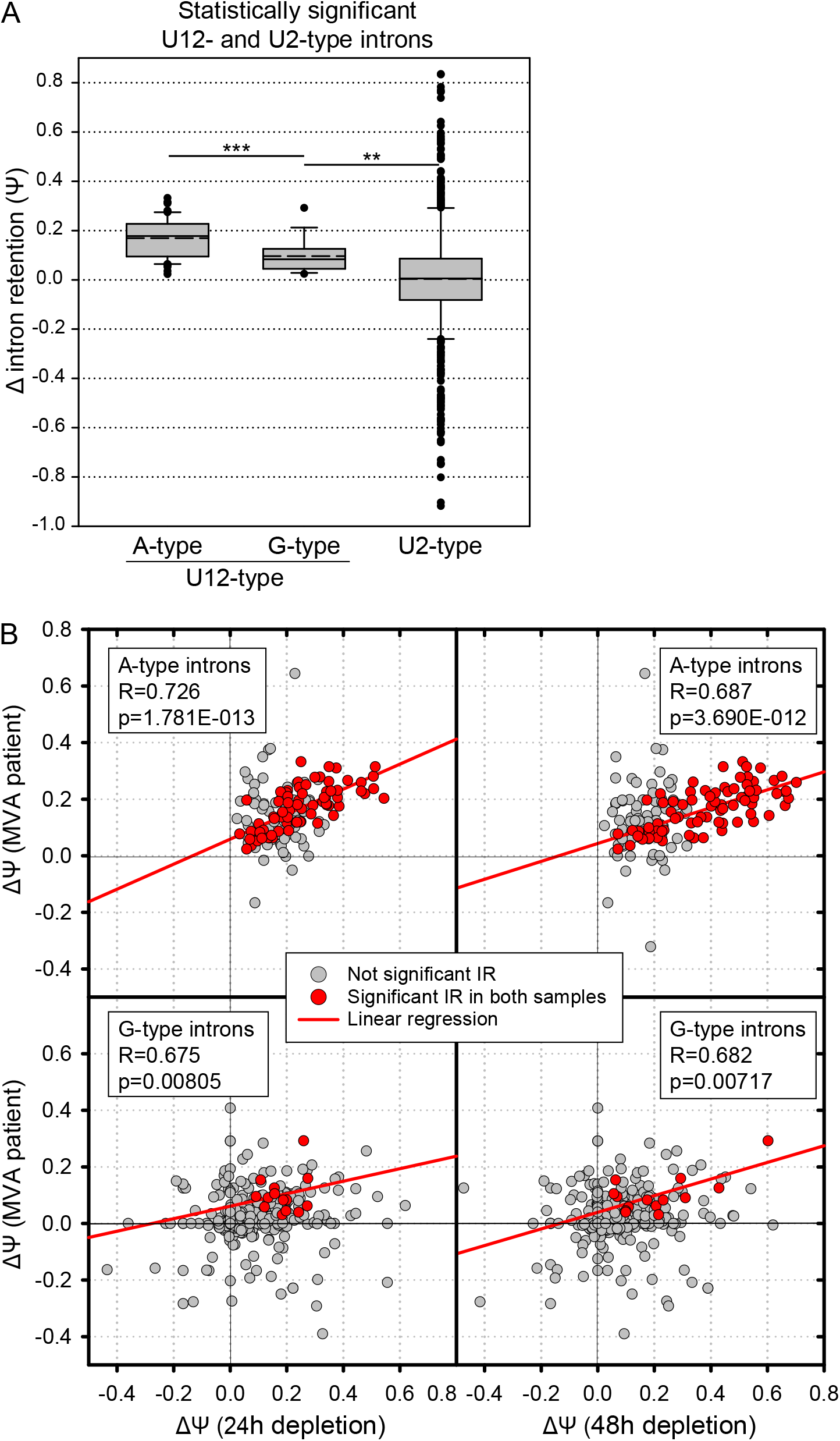
Analysis of the MVA patient intron retention dataset. **A)** Comparison of statistically significant U12 A-type introns (n= 81), U12 G-type introns (n=15), and U2-type introns (n=699). Only introns with on average at least 5 intron-mapping reads were used in the analysis. The boundaries of the boxes indicate 25th and 75th percentiles. Whiskers indicate the 90th and 10th percentiles. Median is indicated with solid line, mean with dashed line inside the box. *** - P<0.001, ** - P<0.01, ns - P>0.05 (Mann-Whitney Rank Sum Test). **B)** Pairwise comparison of minor intron retention values (ΔΨ) between the MVA datset (y-axis) and the CENTAC depletion datasets (x-axis). Red circles indicate introns with statistically significant intron retention values, while with gray circles the retention values were not statistically significant. For the correlation coefficient calculations only the statistically significant intron retention values (red dots) were used.

**Fig. S14.**
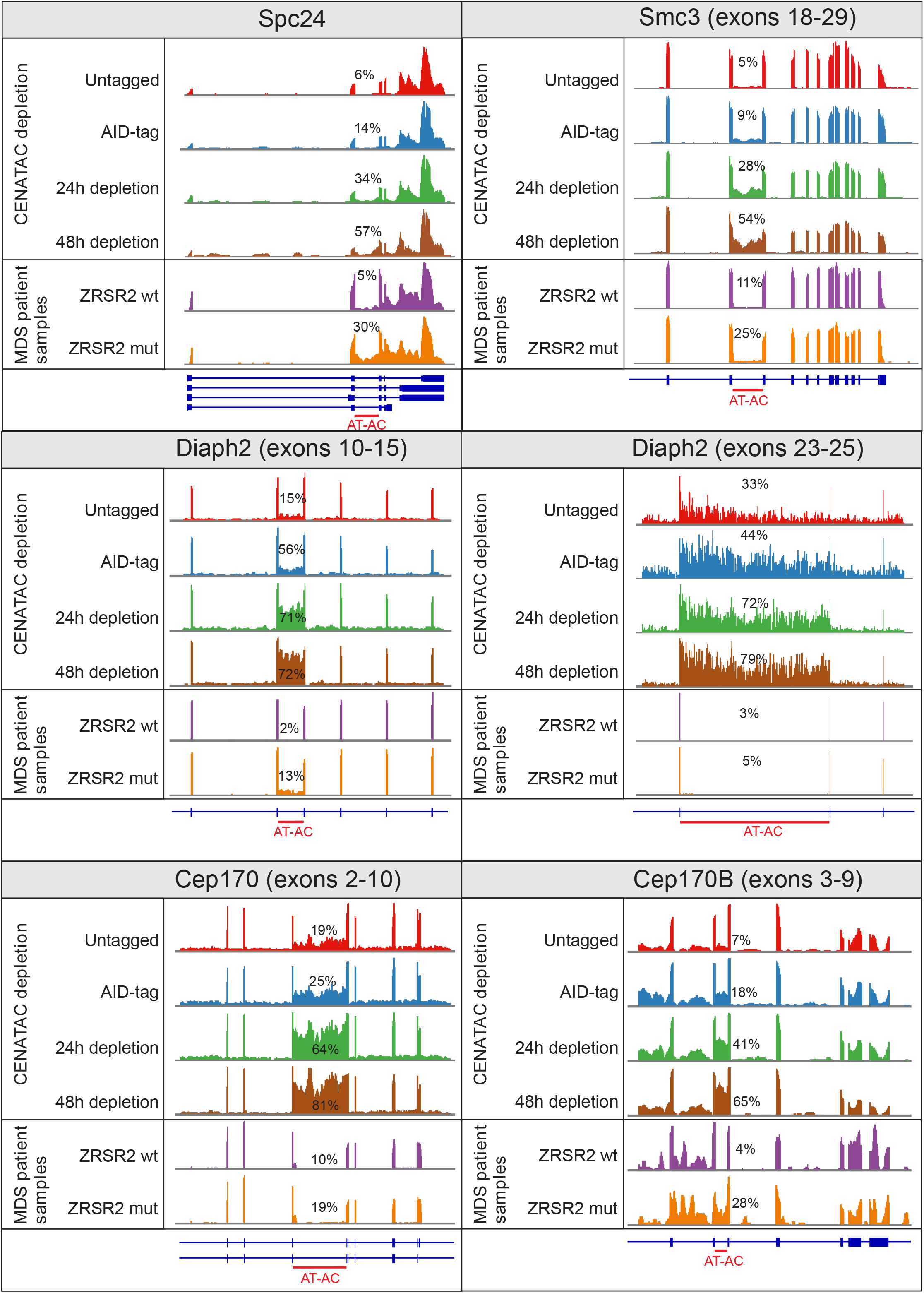
Sashimi plots of mitotic regulators showing the effect of CENATAC depletion in EGFP-AID-CENATAC HeLa cells (24 or 48 hours) or *ZRSR2* mutations in the MDS dataset on AT-AC intron retention (percentages). These genes were manually selected from the genes that were significantly affected by CENATAC depletion (Table S3) with the GO term kinetochore or mitotic spindle. Untagged represents the parental unedited HeLa cell line, AID-tag represents the EGFP-AID-CENATAC HeLa cell line; both were depleted of GAPDH for 48h.

**Fig. S15.**
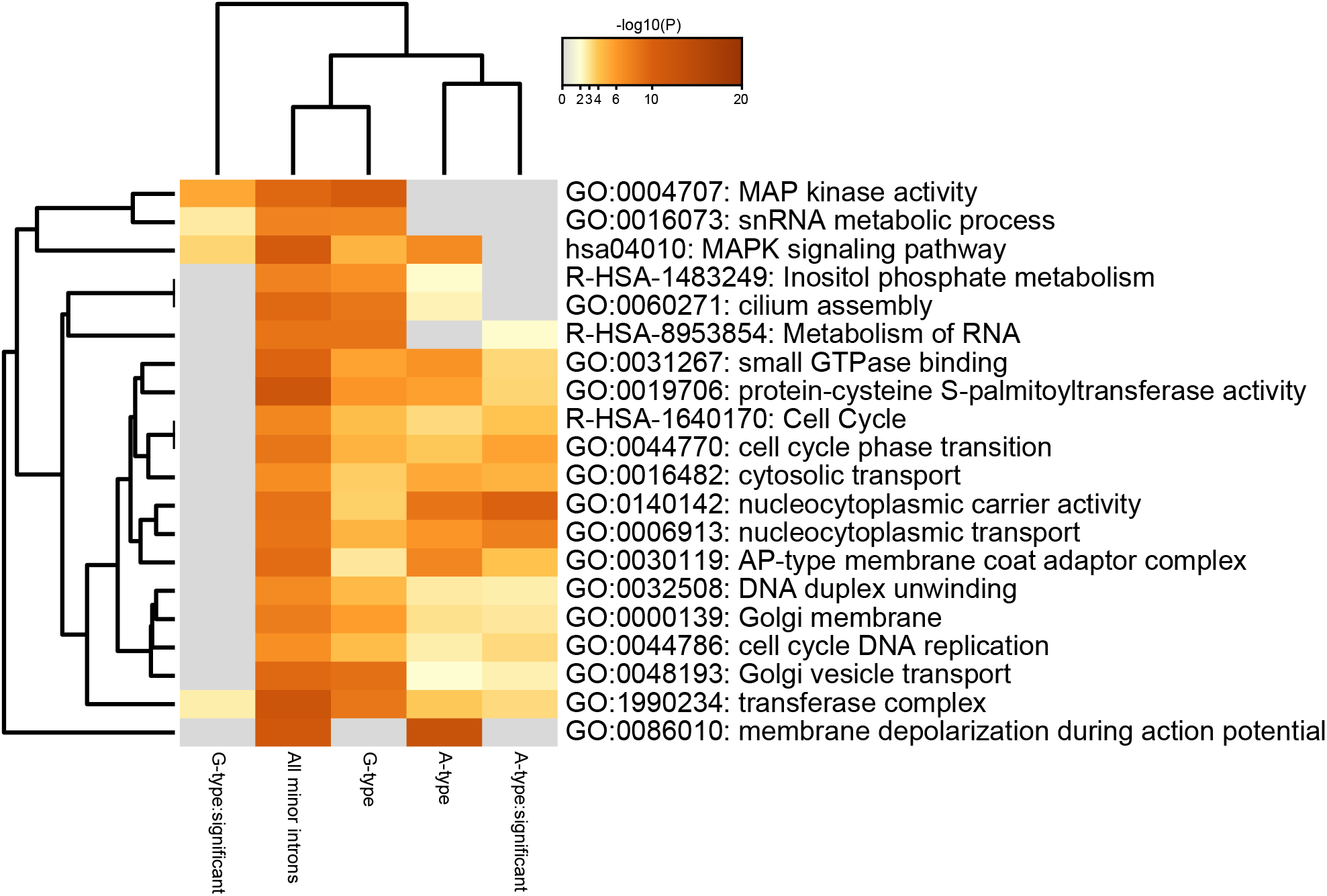
Metascape analysis (metascape.org) of genes containing significantly retained minor introns following 48h CENTAC depletion. Genes with significantly retained A- and G-type introns were analyzed separately (A-type:significant and G-type:significant, respectively) and compared to all genes containing A- or G-type introns (A-type and G-type, respectively) and to all genes containing minor introns. Default parameters were used in the analysis. The color scheme indicates the p-values. Top 20 categories are shows. Table S7 includes the numeric p-values related to the figure. Table S8 includes annotations at the gene level and details of the enriched clusters.

**Fig. S16.**
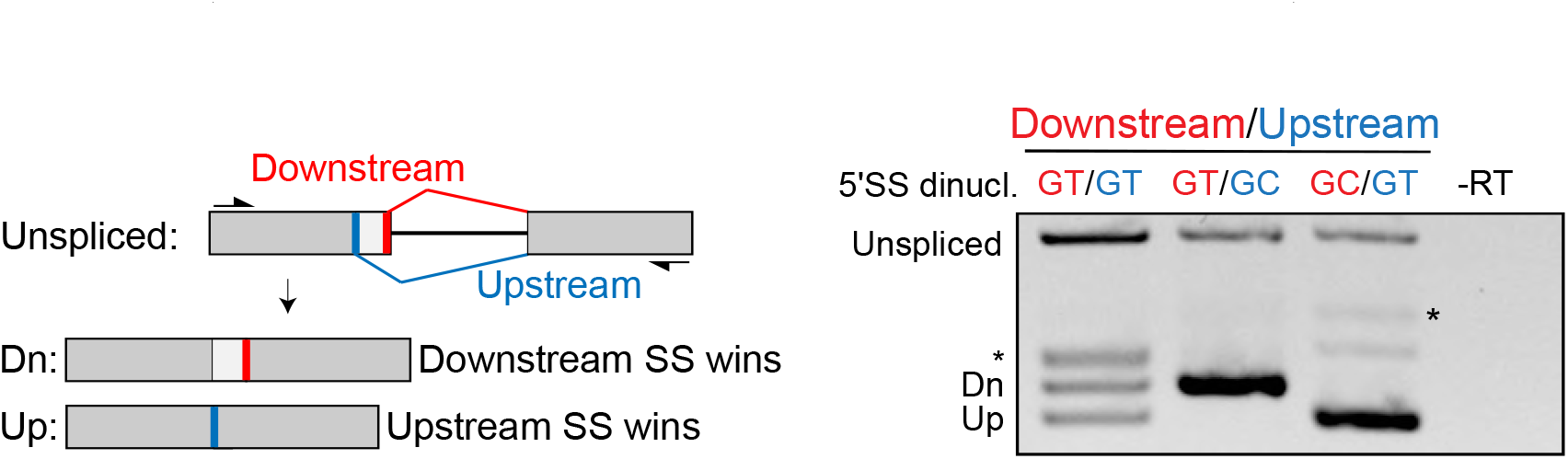
RT-PCR P120 reporter assay to measure the relative usage of GT-AG and GC-AG G-type U12 splice sites in direct competition. Left: schematic diagram showing the overall architecture of the reporter construct with its down- and upstream splice site (thick red and blue bars, respectively) and the products created by splicing (Dn and Up, respectively). Right: RT-PCRs of the reporter with GT-AG or GC-AG splice sites in the down- or upstream positions as indicated above the gel. SS, splice site. 5’SS dinucl., 5’SS dinucleotides. *, PCR product after use of a cryptic U2-type splice site (not shown in the schematic).

## Footnotes

i CENATAC was named after AT-AC introns which make up 84% of U12 A-type introns

